# A basic model of calcium homeostasis in non-excitable cells

**DOI:** 10.1101/2022.12.28.522077

**Authors:** Christina H. Selstø, Peter Ruoff

**Affiliations:** Department of Chemistry, Bioscience, and Environmental Engineering, University of Stavanger, Stavanger, Norway

## Abstract

The level of cytosolic calcium (Ca^2+^) in cells is tightly regulated to about 100 nM (pCa ≈ 7). Due to external stimuli, the basal cytosolic Ca^2+^ level can temporarily be raised to much higher values. The resulting Ca^2+^ transients take part in cell-intrinsic signals, which result in cellular responses. Because of its signaling importance and that high levels of Ca^2+^ can lead to apoptosis, regulation and homeostatic control of cytosolic Ca^2+^ is essential. Based on experimentally known molecular interactions and kinetic data together with control theoretic concepts (integral feedback) we developed a basic computational model describing robust cytosolic Ca^2+^ homeostasis. The aim of the model is to describe the integrative mechanisms involved in cytosolic Ca^2+^ homeostasis in non-excitable cells. From a model perspective, the cytosolic steady state value (set point) of 100 nM is determined by negative feedback loops (outflow controllers), one of these represented by the plasma membrane Ca^2+^ ATPase (PMCA) - calmodulin (CaM) pump and its activation by cytosolic Ca^2+^. Hysteretic behaviors of the Ca pumps and transporters have been added leading to improved kinetic behaviors indicating that hysteretic properties of the Ca^2+^ pumps appear important how cytosolic Ca^2+^ transients are formed. Supported by experimental data the model contains new findings that the activation of the inositol 1,4,5,-tris-phosphate receptor by cytosolic Ca^2+^ has a cooperativity of 1, while increased Ca^2+^ leads to a pronounced inhibition with a cooperativity of 2. The model further suggests that the capacitative inflow of Ca^2+^ into the cytosol at low Ca^2+^ storage levels in the ER undergoes a successive change in the cooperativity of the Store Operated calcium Channel (SOCC) as Ca^2+^ levels in the ER change. Integrating these aspects the model can show sustained oscillations with period lengths between 2 seconds and 30 hours.

**Author Summary:** Cytosolic calcium is subject to a general homeostatic regulation to about 100 nM against a ten thousand times larger extracellular calcium concentration. We investigated the conditions for robust cytosolic and luminal (endoplasmatic reticulum, ER) calcium homeostasis in non-excitable blood and epithelial cells and how external and internal calcium perturbations affect these homeostatic mechanisms. We found that gradual time-dependent (hysteretic) changes of calcium pumps and transporters and their associated cooperativities play an essential role in observed kinetics of the calcium flow in and out of the ER. Using a two-site calcium binding model we quantitatively describe the cytosolic calcium-induced calcium transport out of the ER with a cooperativity of 1, and its inhibition at higher cytosolic calcium concentrations with a cooperativity of 2. For the capacitative Ca entry by Store Operated Calcium Channels (SOCCs) when ER calcium needs to be refilled we find excellent agreement between experimental kinetic data and the model when the cooperativity of luminal calcium changes from 1.3 at 500 *μ*M to 0.8 at 20 *μ*M. Integrating these different aspects of cytosolic and store calcium regulation leads to a basic model for cellular calcium homeostasis, which can show oscillations with period lenths from a few seconds up to 30 hours!

## Introduction

Calcium is one of the most abundant and versatile cations in organisms and the human body. Not only important for the maintenance of the skeleton, calcium is also important for the overall health and signaling processes [1]. As a second messenger calcium plays a role in nearly all physiological processes ranging from fertilization, photoreceptor regulation to cell death [1–6].

The concentration of free cytosolic Ca^2+^ in a resting cell is kept at around 100 nM while in the extracellular environment the Ca^2+^ concentration is about ten thousand times higher, i.e., about 1 mM. To keep the level of cytosolic Ca^2+^ robustly at such low levels any perturbation in cytosolic Ca^2+^ is opposed by compensatory homeostatic mechanisms [7]. These mechanisms consist of negative feedback loops where transporters remove excess of cytosolic Ca^2+^ by excreting it or moving it to cellular stores.

An overview of the components involved in cytosolic Ca^2+^ homeostasis is shown in Fig 1. In non-excitable cells, the Arachidonic Acid Regulated Ca^2+^ Channel (ARCC) and the Store Operated Ca^2+^ Channel (SOCC) constitute the main Ca^2+^ entry pathways into the cell. They are both Orai channels connecting to STIM in either the plasma membrane (PM) or the ER membrane for ARCC and SOCC respectively [8]. Orai proteins are proteins found in the plasma membrane which constitute the pore subunit of the channels [8, 9]. The difference between these channels is that while SOCCs are dependent on the Ca^2+^ level in the ER store, the ARCCs are not activated by Ca^2+^ in the ER, but by extracellular signals. This occurs by first activating arachidonic acid, which in turn will then activate the ARCCs. While STIM is assumed to be bound to ARCC, ARCC is not explicitly included in the model. The SOCCs, also known as Calcium Release Activated Channels (CRACs), are responsible for the inflow of Ca^2+^ into the cell to refill the ER with calcium, when depleted by leaks [10] or by signaling events [8, 11, 12].

**Fig 1.**
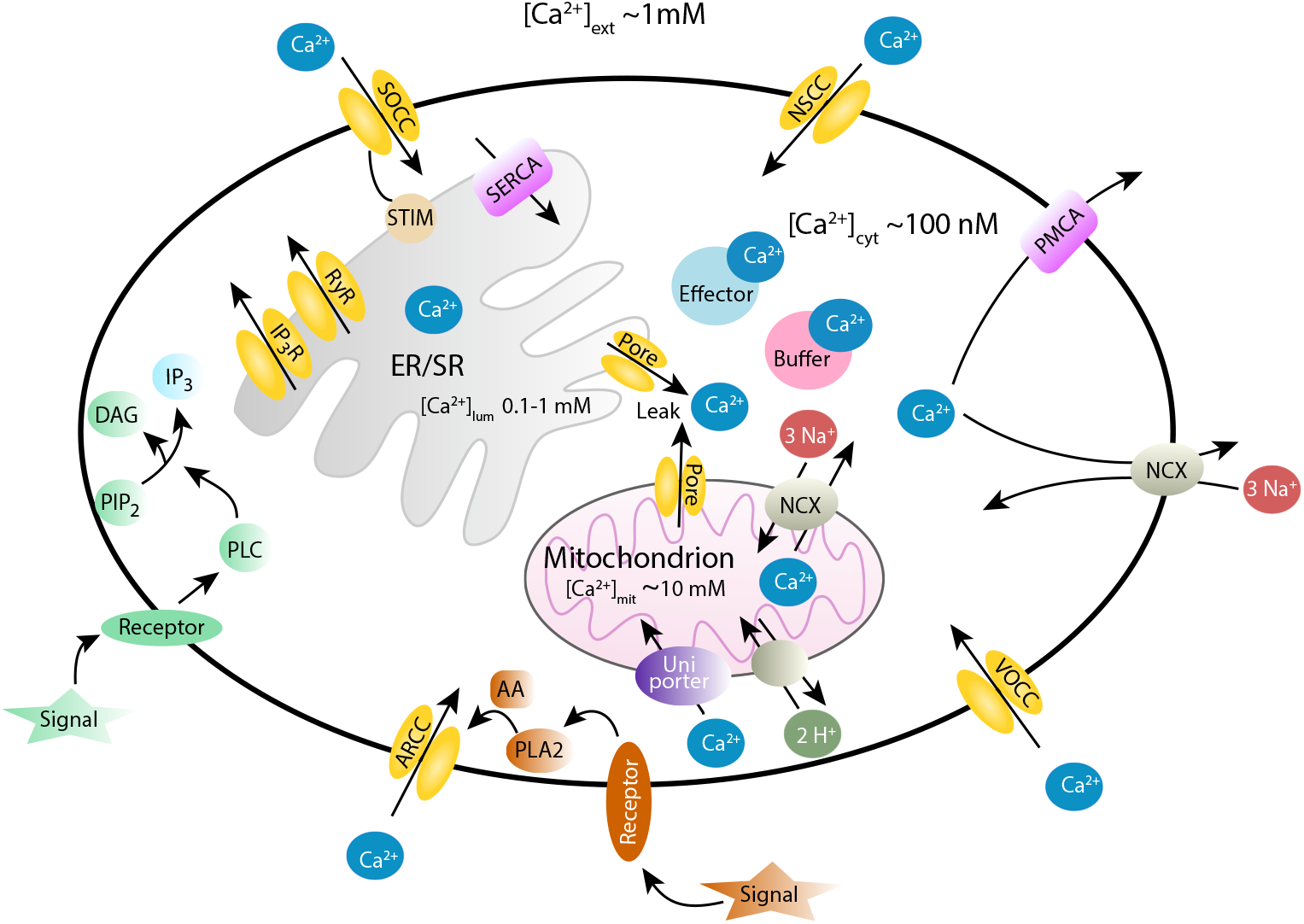
Overview of Ca^2+^-ion transport in an eukaryotic cell. The illustration shows the channels, pumps, receptors, and organelles involved in the Ca^2+^ transport to and from the cytosol. The channels for Ca^2+^ influx from the extracellular space include the Voltage Operated Calcium Channels (VOCCs) in excitable cells, ligand operated Ca^2+^ channels like the Arachidonic acid Regulated Calcium Channels (ARCCs), Store Operated Calcium Channels (SOCCs) and Non-Specific Cation Channels (NSCC). Ca^2+^ can also enter the cytosol from organelle stores such as the endoplasmic reticulum (ER) and mitochondria. The Inositol 4,5-Trisphosphate Receptor (IP_3_R) and Ryanodine Receptors (RYRs) release Ca^2+^ into the cytosol in a Ca^2+^ dependent manner. There is also evidence that there is a Ca^2+^ leakage from the ER or organelle stores into the cytosol. The IP_3_R channel is activated by IP_3_. IP_3_ is the result of a signal transduction chain originating from a signal which activates a G-protein coupled receptor and produces phospholipase C (PLC). PLC catalyzes the hydrolysis of phosphatidylinositol 4,5-bisphosphate (PIP_2_) which produces IP_3_. Mitochondria can expel Ca^2+^ through pores and ion exchangers (Na^+^/Ca^2+^ and H^+^/Ca^2+^). Plasma Membrane Ca^2+^ ATPase (PMCA) and the sodium-calcium exchanger (NCX) transport cytosolic Ca^2+^ into the extracellular space. The Sarco/Endoplasmic Reticulum Calcium ATPase (SERCA) is responsible for the Ca^2+^ transport into the ER, while transport of Ca^2+^ into mitochondria occurs via an uniporter. In addition, proteins both present in the cytosol and organelles can bind Ca^2+^ and act as buffers (buffer proteins), like calsequestrin or calreticulin, while other Ca^2+^ binding proteins, such as calmodulin (CaM), mediate Ca^2+^ effects (effector proteins). For references about these processes see main text.

When Ca^2+^ enters the cytosol through channels in the plasma membrane (PM), the cytosolic concentration of Ca^2+^ is regulated to about 100 nM by binding to buffering proteins, by organelle sequestration of Ca^2+^ and by extruding Ca^2+^ through the PM. There are several Ca^2+^ binding proteins which act either as buffers (buffer proteins) or mediate an effect by Ca^2+^ (effector proteins). Calmodulin (CaM) is an important effector protein which takes part in the Ca^2+^ activation of PMCA and NCX, but plays also an important role in Ca^2+^ dependent signaling [13, 14]. The PMCA appears to be the most important efflux path for Ca^2+^ in non-excitable cells, although some non-excitable cells have in addition the Na^+^/Ca^2+^ exchanger (NCX). Some controversy exists on how important the role of NCX is in non-excitable cells, whereas in excitable cells NCX appears most important for removing Ca^2+^ from the cytosol through the PM [15, 16]. The PMCA has been thought of having a housekeeping function, because of its high affinity to Ca^2+^ and a relatively low throughput thereby bringing cytosolic Ca^2+^ concentrations down to their resting levels. The PMCA comes in different isoforms with various Ca^2+^ affinities and transport rates, which have been suggested as a fine tuner of signals in localized areas of the cell [15, 17, 18]. NCX, on the other hand, is a low affinity transporter with a high throughput in Ca^2+^ removal, which is needed in excitable cells where rapid increases in cytosolic Ca^2+^ concentrations occur.

In contrast to the cytosol, both the ER and mitochondria contain much higher Ca^2+^ concentrations, i.e., 0.1 - 1mM in the ER, and around 10mM in mitochondria. In this work, the mitochondria being involved in the storage and buffering of Ca^2+^ are not included in the model. On the other hand, the ER, which holds important roles in both signaling, in forming Ca^2+^ transients, and in Ca^2+^ storage, has been included as part of the cytosolic and organelle Ca^2+^ homeostatic machinery. As mentioned, SOCCs are mainly responsible for refilling the ER with Ca^2+^ by an ATPase called the sarco/endoplasmic reticulum Ca^2+^ ATPase (SERCA), when Ca^2+^ is depleted in the ER by either the inositol 4,5-trisphosphate receptors (IP_3_R) or by ryanodine receptors (RyR), both located in the ER membrane. Since the RyR’s are mainly responsible for the ER-Ca^2+^ depletion in excitable cells, the IP_3_R channel is considered in the model as the only Ca^2+^ outflow path from the ER, in addition to leakage [10]. IP_3_R is activated both by IP_3_ and by cytosolic Ca^2+^ [19]. The activation and inhibition kinetics of IP_3_R are described in more detail below.

## Materials and methods

Rate equations were solved by using the Fortran subroutine LSODE [20]. When not mentioned otherwise, concentrations and time units are in *μ*M and seconds (s), respectively. Plots were generated with gnuplot (www.gnuplot.info) and edited with Adobe Illustrator (adobe.com). Experimental data were extracted from graphs by using GraphClick (https://graphclick.en.softonic.com/mac). The analyses of experimental data were done with gnuplot or Excel, and then implemented into the model. The program Cn3D [21] was used for the structural analysis of rat IP_3_R [22]. To make annotations simpler, concentrations of compounds are denoted by compound names without square brackets. The Supporting Informations contain source files, compiled binaries, and instructions how to execute the different models.

### Illustrating integral control: A PMCA-based minimal model

The role of Ca^2+^ in signaling is complex and so are the Ca^2+^ fluxes and transport paths in the cell. The intention of this work has been to get an understanding of how robustness in cytosolic Ca^2+^ homeostasis can be achieved while allowing the occurrence of experimentally observed Ca^2+^ transients during an inflow perturbation of Ca^2+^ into the cytosol. The model was developed by starting initially with a simple set of regulatory elements where experimentally known pathways and dynamic properties were then successively added. The initial (minimal) model contains the Ca^2+^ pump PMCA as the essential regulatory element together with Ca^2+^-binding buffer proteins (lumped into variable B) and the Ca^2+^-binding effector protein Calmodulin (variable M). Perturbation of cytosolic Ca^2+^ occurs by a constant inflow *k*_1_ of external Ca^2+^ into the cytosol.

To achieve cytosolic Ca^2+^ homeostasis the PMCA is considered to be part of a negative feedback loop in maintaining a low and stable cytosolic Ca^2+^ concentration. In addition we include integral control in the loop (Fig 2A), which is a concept from control engineering [23–32]. Integral control (or integral feedback) allows to maintain robust homeostasis, i.e. keeping in our case the cytosolic Ca^2+^ level at a given set-point 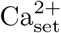 for different but constant (*k*_1_) inflow perturbations. In general, integral control is achieved to integrate with respect to time the error *ϵ*, which is the difference between the Ca^2+^ set-point 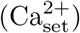 and the actual cytosolic Ca^2+^ concentration. The integrated error is then used to compensate for changes in the cytosolic Ca^2+^ concentration (Fig 2A).

**Fig 2.**
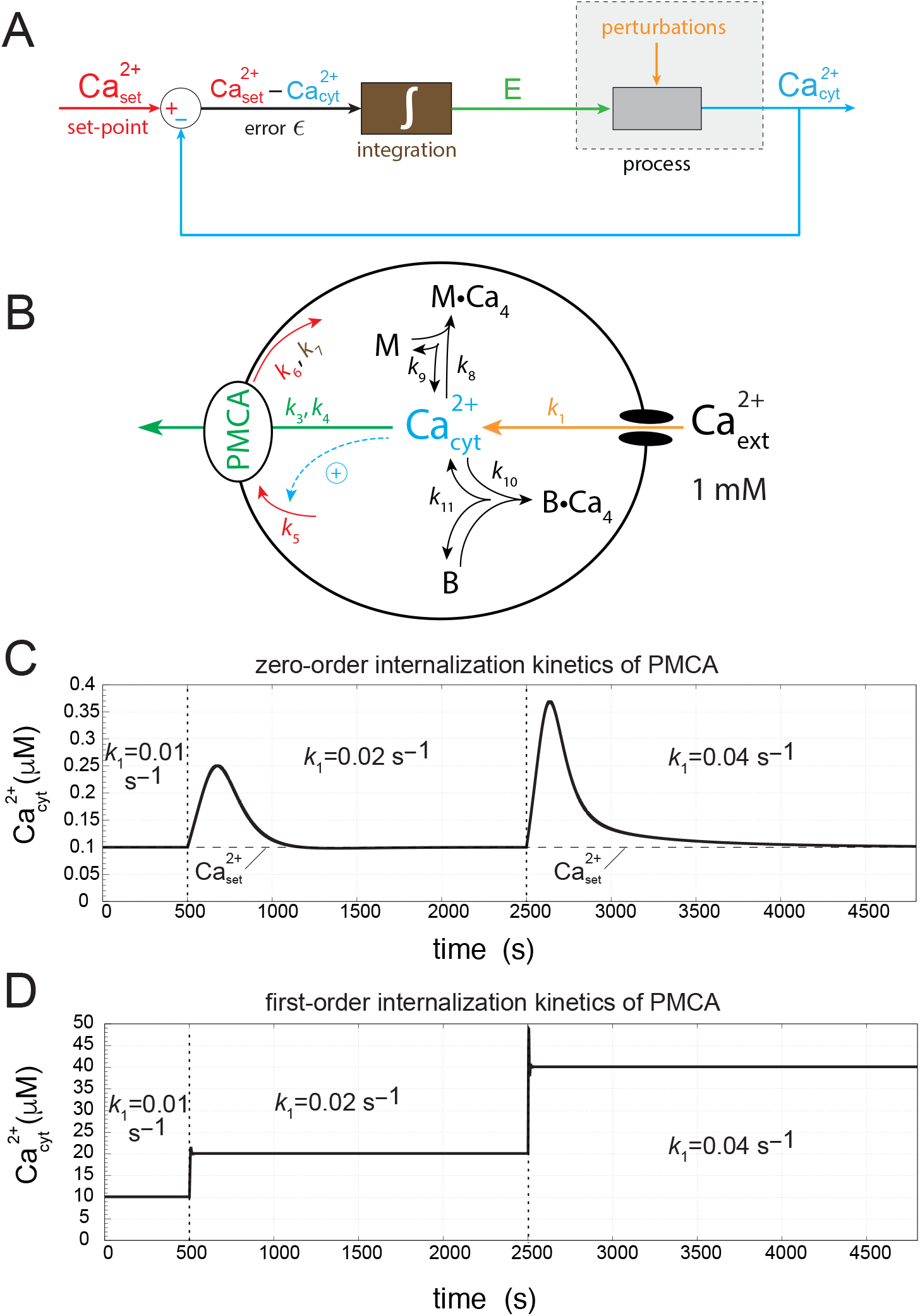
Minimal feedback model of cytosolic Ca^2+^ homeostasis with and without integral control. (A) General flow diagram of negative feedback with integral control. The red color indicates the set-point of the controlled variable 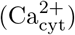 while the blue color refers to its actual value. Integration (brown color) of the error (difference between set-point and actual value) leads to the integrated error *E* (outlined in green) which for step-wise inflow perturbations *k*_1_ (orange color) will precisely move the controlled variable (blue) to its set-point [24]. (B) Minimal molecular model of PMCA-based cytosolic Ca^2+^ homeostasis with color codes matching the flow diagram in panel (A). M and B denote two Ca^2+^ binding proteins with arbitrary binding properties. The kinetics of PMCA internalization (indicated by rate constants *k*_6_ and *k*_7_) determines whether integral control is invoked or not. (C) Zero-order PMCA internalization kinetics leads to integral control and robust homeostasis. In this case the 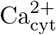 concentration returns to the set-point 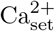 for the different inflow perturbation rates *k*_1_. (D) Loss of robust 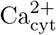 homeostasis when PMCA internalization kinetics is no longer zero-order but first-order. For details, see main text and Supporting Information S1 Program.

There are presently three main kinetic approaches how integral control can be achieved in a chemical system. One approach, which we use here, is based on a zero-order removal of the negative feedback species (controller species) [23, 26, 28]. The second approach (antithetic control) is using two controller species, which react with each other either directly [30, 31] or via an enzyme [33], while one of the controllers feeds back negatively to the controlled variable. In the third approach the controller variable is produced by first-order autocatalysis, but needs to be removed by a first-order reaction [34–36].

Fig 2B shows a molecular representation of Fig 2A using a zero-order removal of the controller variable PMCA (our initial model). The following PMCA-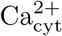 negative feedback loop is considered: cytosolic Ca^2+^ is activating PMCA [37] and thereby transporting 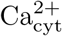 out of the cell and opposing the effect of an increasing 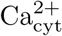. PMCA inactivation is considered to occur by internalization [38] as for other plasma membrane proteins [39]. Activation and deactivation of PMCA are outlined in red, and as we will show below are important for determining the Ca^2+^ set-point in the cytosol. To achieve integral control with a robust set-point we focus here on a zero-order internalization of PMCA as a requirement for integral control.

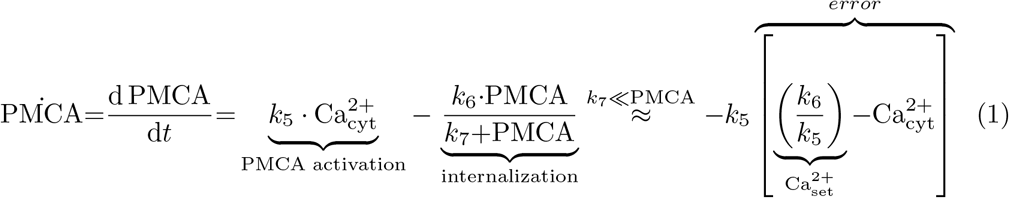

Zero-order kinetics for PMCA internalization is obtained when *k*_7_ ≪ PMCA, i.e. PMCA binds tightly to the enzymatic internalization machinery where *k*_7_ plays the role of a dissociation/Michaelis constant. Eq. 1 shows the rate equation for PMCA, its activation and internalization terms and the reorganization of the equation into the error term between set-point and the actual Ca^2+^ concentration when zero-order condition applies. Fig 2C shows the robust perfect adaptation of cytosolic calcium to its set-point for different *k*_1_ inflow rates when zero-order internalization kinetics are applied.

On the other hand, when internalization of PMCA becomes first-order with respect to PMCA, i.e., when

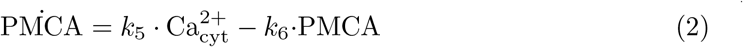

then the robustness of the PMCA negative feedback loop is lost and 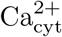 increases proportionally with *k*_1_ (Fig 2D). For a complete set of the rate equations, rate constant values and initial concentrations, see Supporting Information S1 Program.

## Results and discussion

### Overview of the cellular model

Fig 3 shows the model considered here, including all parts and dynamic behaviors which we deemed necessary for describing cellular Ca^2+^ homeostasis. As a Ca^2+^ store we have focused on the endoplasmatic reticulum (ER), but have not included other organelles such as mitochondria.

**Fig 3.**
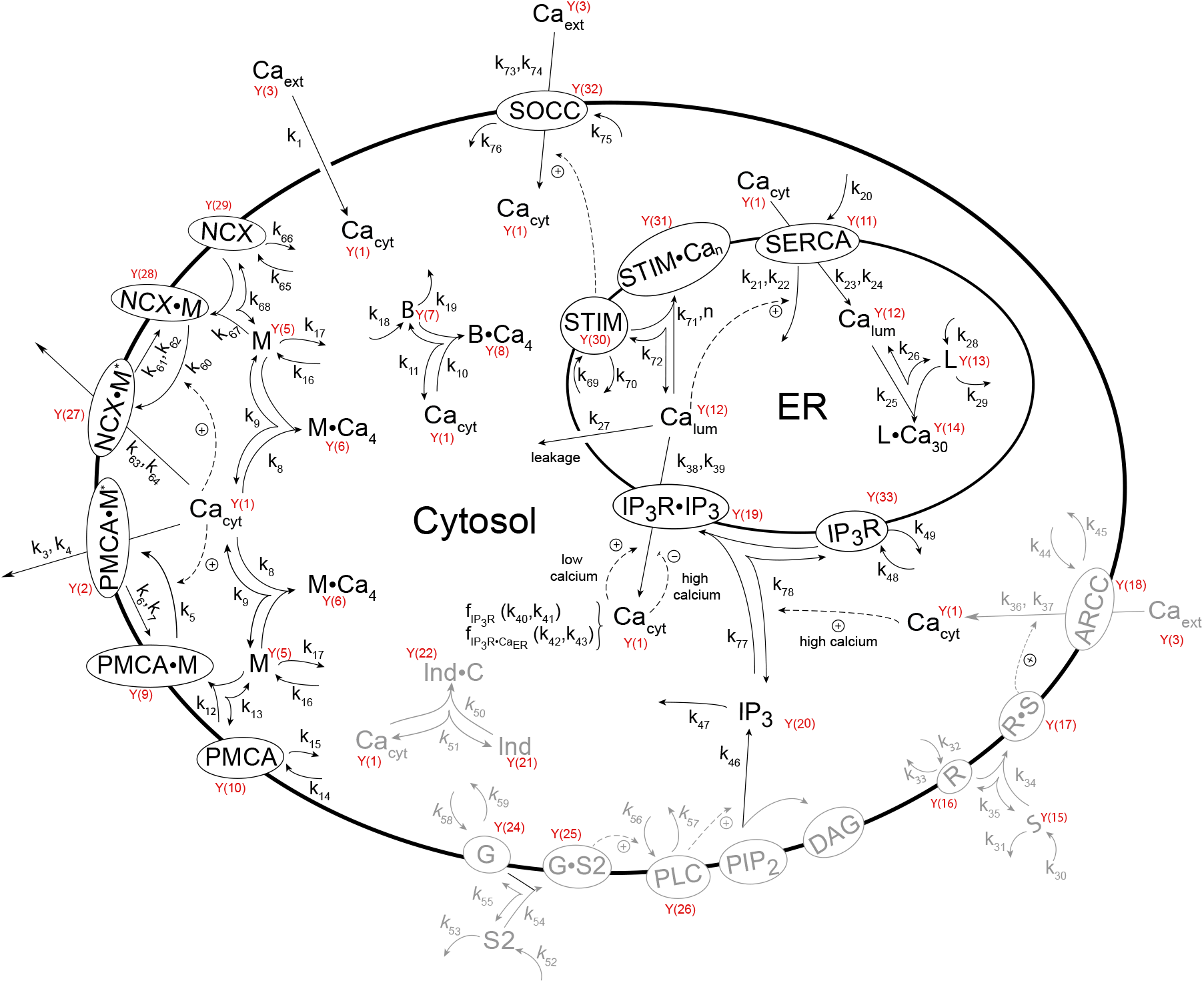
Model of Ca^2+^ homeostasis in the cytosol and endoplasmatic reticulum (ER). Components and rate constants of the model are shown in this overview figure. The Y(i)’s outlined in red are the LSODE Y vector’s components [20], which are assigned to each biochemical species in the model. Arrows represent reactions of the involved species while dashed lines indicate signaling pathways, i.e., how components activate or inhibit each other. The grayed-out reactions with rate constants and Y(i)’s are included in the model, but are not explicitly presented. The set of rate equations together with a list of abbreviations and the LSODE Y(i) assignments are described in S1 Text.

While the primary focus of our work was on Ca-homeostasis and not on oscillations, we observed that negative feedback loops have the potential to oscillate, either regular or chaotic, and that even under such conditions the negative feedback mechanisms are able to maintain robust homeostasis [40, 41]. It was therefore not surprising that the model can relatively easy exhibit oscillations and still maintain Ca-homeostasis in both the cytosol and in the ER. At the end of the paper we briefly describe some of these oscillations, which, dependent on parameter values can have period lengths from a few seconds up to 30 hours!

In the following we describe in more detail the different aspects of the model in comparison with experimental results.

### The plasma membrane Ca^2+^ ATPase

The cell type we consider first are erythrocytes. They lack most organelles including the nucleus, mitochondria, the Golgi apparatus and the endoplasmatic reticulum (ER). Ca^2+^ homeostasis in these cells is maintained by PMCA, which appears to be the most abundant pump in these cells. To transport one Ca^2+^ out of the cell the PMCA requires one ATP and exchanges H^+^ ions for Ca^2+^. The stoichiometry of the Ca^2+^/H^+^ exchange is controversial [15], but neither ATP nor H^+^ ions are explicitly included in the model (which have their own homeostasis mechanisms). Maximum activation of the PMCA pump requires the presence of calmodulin (M); its importance in PMCA activation is widely agreed upon [37, 42–44]. PMCA can also function and be partially active without calmodulin, probably due to acidic phospholipids, but only to about half of its maximal activity [37, 43].

Fig 4 shows part of the model with calmodulin (M) activating PMCA by forming PMCA•M. The PMCA•M complex is further activated by Ca^2+^ leading to PMCA•M^*^, which is the form with the maximum Ca^2+^ pump activity. PMCA is a high affinity pump with a K_d_ (with respect to Ca^2+^) of 0.1–1 *μ*M when calmodulin is bound, but a K_d_ of about 10–20 *μ*M when calmodulin is absent [15, 45].

**Fig 4.**
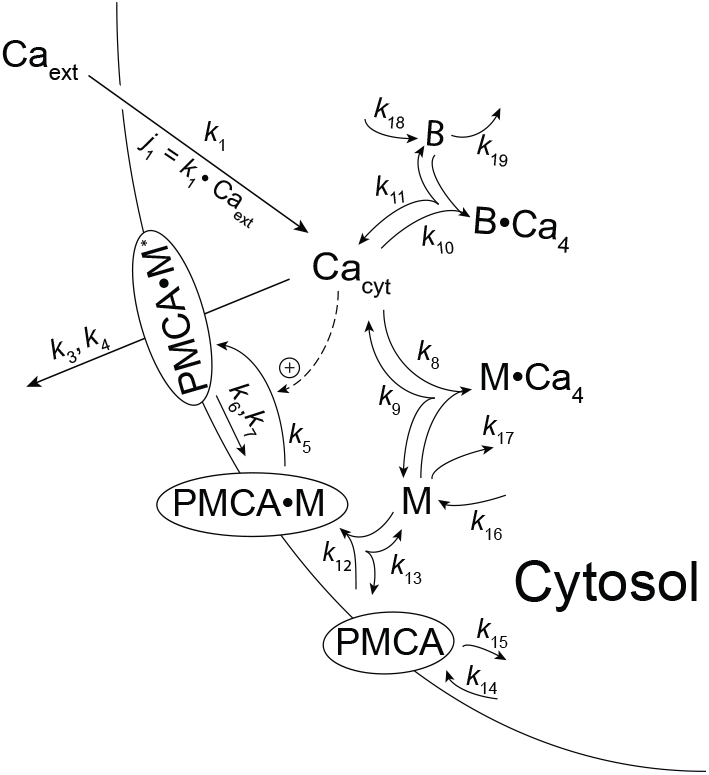
Calmodulin and Ca^2+^ activation of PMCA. The figure shows the activation of PMCA by calmodulin (M) and Ca^2+^. Although experiments have shown that PMCA can apparently transport Ca^2+^ in the absence of calmodulin, but less effectively [15, 45], we have in the model not included a PMCA based Ca^2+^ transport without calmodulin.

In our calculations we have taken the set-point for cytosolic Ca^2+^ to be 0.1 *μ*M. To achieve robust homeostasis we consider, as for the minimal model (Fig 2), a first-order activation of PMCA by Ca^2+^ with rate constant *k*_5_ (Fig 4) leading to the active form PMCA•M^*^. The inactivation of PMCA•M^*^ is considered a zero-order process with respect to PMCA•M^*^. The set-point for cytosolic Ca^2+^ is given by the steady state condition for PMCA•M^*^, i.e.,

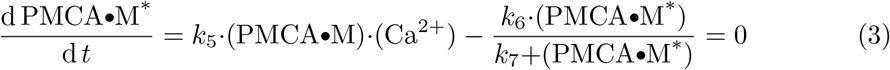

where *k*_7_ plays the role of a dissociation (Michaelis) constant. Applying zero-order kinetics with respect to PMCA•M^*^, *k*_7_ ≪ *PMCA*•*M* ^*^, the steady state condition of Eq. 3 becomes

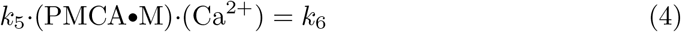

which determines the steady state value of cytosolic Ca^2+^

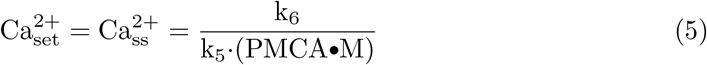

and which is independent of the Ca inflow perturbation *j*_1_=*k*_1_·Ca_ext_ (see Fig 4).

Since we applied first-order kinetics with respect to PMCA•M in its Ca^2+^ activation, the set-point of Ca^2+^ depends on the concentration of PMCA•M. This leads to the possibility of a *variable* set-point, as has been argued for by Mrosovsky on general grounds [46]. As will be shown/discussed below such a variable set-point does well describe the experimental results.

### PMCA kinetic parameters

In the following we discuss the parameter values used in the model focusing first on red blood cells. To the extent this is possible we try to use or derive rate and bindings constants primarily from *in vivo* studies.

For PMCA’s pump rate *v*_*pump*_ we are using a simple Michaelis-Menten type expression

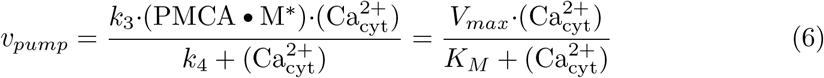

where PMCA•M^*^ denotes the active form of the pump, *V*_*max*_=*k*_3_·(PMCA • M*) with *k*_3_ and *k*_4_ as turnover number (*k*_*cat*_) and dissociation (Michaelis) constant, respectively.

#### PMCA K_M_ values

The enzyme data base BRENDA [47] has 6 *K*_*M*_ entries for human PMCA and SERCA (Ca-ATPases) with values for erythrocytes, liver, and heart ranging from 0.013–20 *μ*M [48–50]. For red blood cells, we have analyzed the *in vitro* data by Niggli et al. [51], which indicate a *K*_*M*_ value for PMCA of about 1 *μ*M (see S2 Text).

By using two protocols, Kubitscheck et al. [52] determined the *K*_*M*_ value of PMCA in single red blood cells to 24 *μ*M and 1 *μ*M. The 24 *μ*M value has been interpreted by the authors due to an inactive calmodulin in one of their protocols. A *K*_*M*_ value of about 0.2 *μ*M was indicated by Bruce [53] for non-excitable cells, while Blaustein [54] refers to a *K*_*M*_ value of 0.1 *μ*M, but in both cases no explicit reference to experimental work was provided. However, our calculations indicate no practical differences when *K*_*M*_ values of 0.1 *μ*M or 1 *μ*M are used (see S2 Text).

#### PMCA Turnover number and V_max_ values

Entries in BRENDA for the human PMCA turnover number (*k*_*cat*_) vary from 9.5–149 s^−1^. In Bradshaw and Dennis’ “Handbook of Cell Signaling” Blaustein lists a *k*_*cat*_ value of 30 s^−1^ [54] (without reference to experiments), while Chen et al. reported a *k*_*cat*_ of about 200 s^−1^ in rat cochlear hair cells [55].

Considering red blood cells Schatzmann [56] found a *V*_*max*_ for PMCA of 148 *μ*M/min. A similar value of 120 *μ*M/min (average, see S2 Text) was obtained by Dagher and Lew [57]. In the Dagher and Lew experiments, also using erythrocytes, the zero-order kinetic extrusion rate of PMCA was determined by using the ionophore A23187 for massively loading the cells with Ca^2+^ and then inhibiting A23187 by using CoCl_2_ (Fig 5A). Due to the large amount of cytosolic Ca^2+^ in comparison with the pump’s *K*_*M*_ value the pump runs at maximum speed (*V*_*max*_). By using the same method Tiffert et al. [58] measured the PMCA-mediated Ca^2+^ extrusion rate in red cells from freshly drawn blood in relation to the hemoglobin content of the cells. By assuming that one red blood cell contains 270 × 10^6^ hemoglobin molecules and that one red blood cell has a volume of approximately 10^−13^ liter, *V*_*max*_ for the pump was estimated to 234 *μ*M/min (S2 Text for details). In addition, Tiffert et al. [58] alternatively measured the pump rate in relation to the number of cells. Using these values we estimate a *V*_*max*_ of about 156 *μ*M/min (S2 Text) close to the value originally determined by Schatzmann [56].

**Fig 5.**
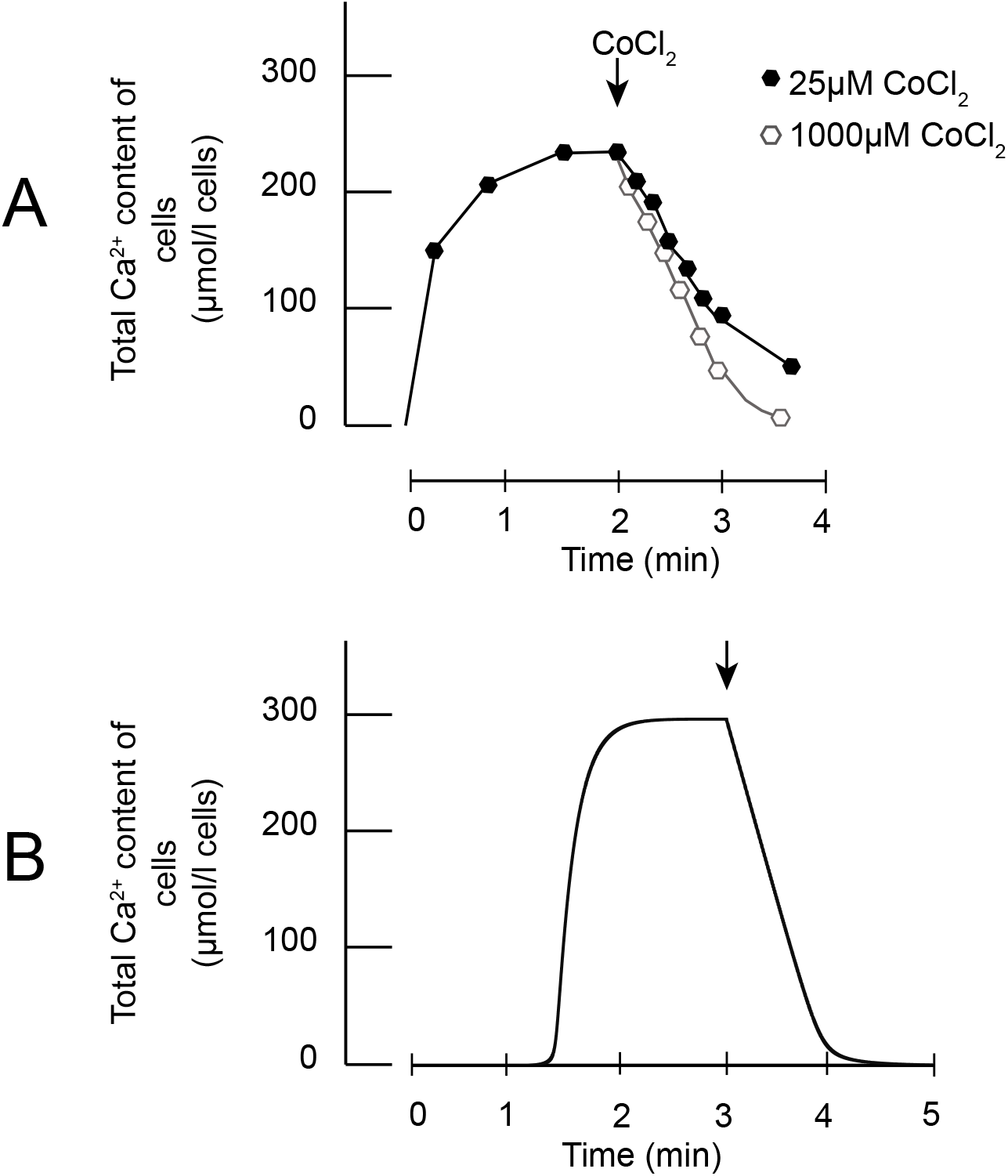
Changes in Ca^2+^ concentration for extrusion rate determination in erythrocytes. Panel A shows experimental plots adapted from Dagher and Lew [57], where the ionophore A23187 is used to let Ca^2+^ enter the cytosol until PMCA (and other pumps) can match the Ca-inflow, which leads to a steady state in cellular Ca^2+^. Then, CoCl_2_ was added to block the ionophore (indicated by arrow). With higher CoCl_2_ concentrations the rate of Ca^2+^ extrusion becomes linear with respect to time and shows close to zero-order kinetics with respect to cellular Ca^2+^. This indicates that the pump’s removal of Ca^2+^ from the cells is at a maximum rate (V_max_). Panel B shows the results from a corresponding model calculation. Rate equations, parameter values and initial concentrations are given in ‘S2 Program’ together with the Fortran source code and executables.

Fig 5A shows the above referred experiment by Dagher and Lew [57] when PMCA (with possibly other pumps) balance a large inflow of Ca^2+^ by use of the ionophore A23187. When the ionophore is blocked the kinetics of the pumps’ Ca^2+^ extrusion can be observed. Fig 5B shows a corresponding set of model calculations.

### PMCA activation and cytosolic calcium profile

Using bovine endothelial cells Sedova and Blatter [59] investigated PMCA activation by following the inflow of Ca^2+^ into the cytosol when 2mM extracellular calcium is applied. The cellular inflow of Ca^2+^ occurs, because the calcium level in the ER is low and leads to a so-called ‘capacitative Ca entry’ [60] into the cell to refill the ER with calcium (see also section ‘Store operated Ca^2+^ entry’ below). The inflow of calcium into the cytosol activates PMCA, which reduces the level of cytosolic Ca^2+^ leading to a biphasic response. Fig 6A (phases 1 and 2) shows the increase and decrease in cytosolic Ca^2+^ concentration when cells are treated with 2 mM extracellular calcium. In phase 3 the extracellular calcium is washed out and cells were treated with the PMCA inhibitor carboxyeosin. Then the inhibitor was washed out and 2mM extracellular calcium was reapplied. Sedova and Blatter now observed a slower increase of cytosolic Ca^2+^ (Fig 6A, phase 4). We used the parameters *k*_1_ (inflow of Ca^2+^ into the cytosol) and *k*_3_ (maximum rate of PMCA) to describe the observed behaviors, but are neglecting here the further transfer of cytosolic calcium into the ER. The slower increase of cytosolic Ca^2+^ in phase 4 is based on two assumptions we made: (i) carboxyeosin partly inhibits the inflow of calcium into the cytosol. This assumption is based on results by Choi and Eisner for rat myocytes [61], who observed that the inflow of Ca^2+^ into the cytoplasm was inhibited by carboxyeosin. The other assumption was that PMCA still remained partly inhibited, as not all carboxyeosin may have been washed out and/or the reactivation of PMCA is slow due to a hysteretic property of the pump. Panels B and C in Fig 6 show the calculated cytosolic Ca^2+^ profiles and the underlaying changes in *k*_1_ and *k*_3_.

**Fig 6.**
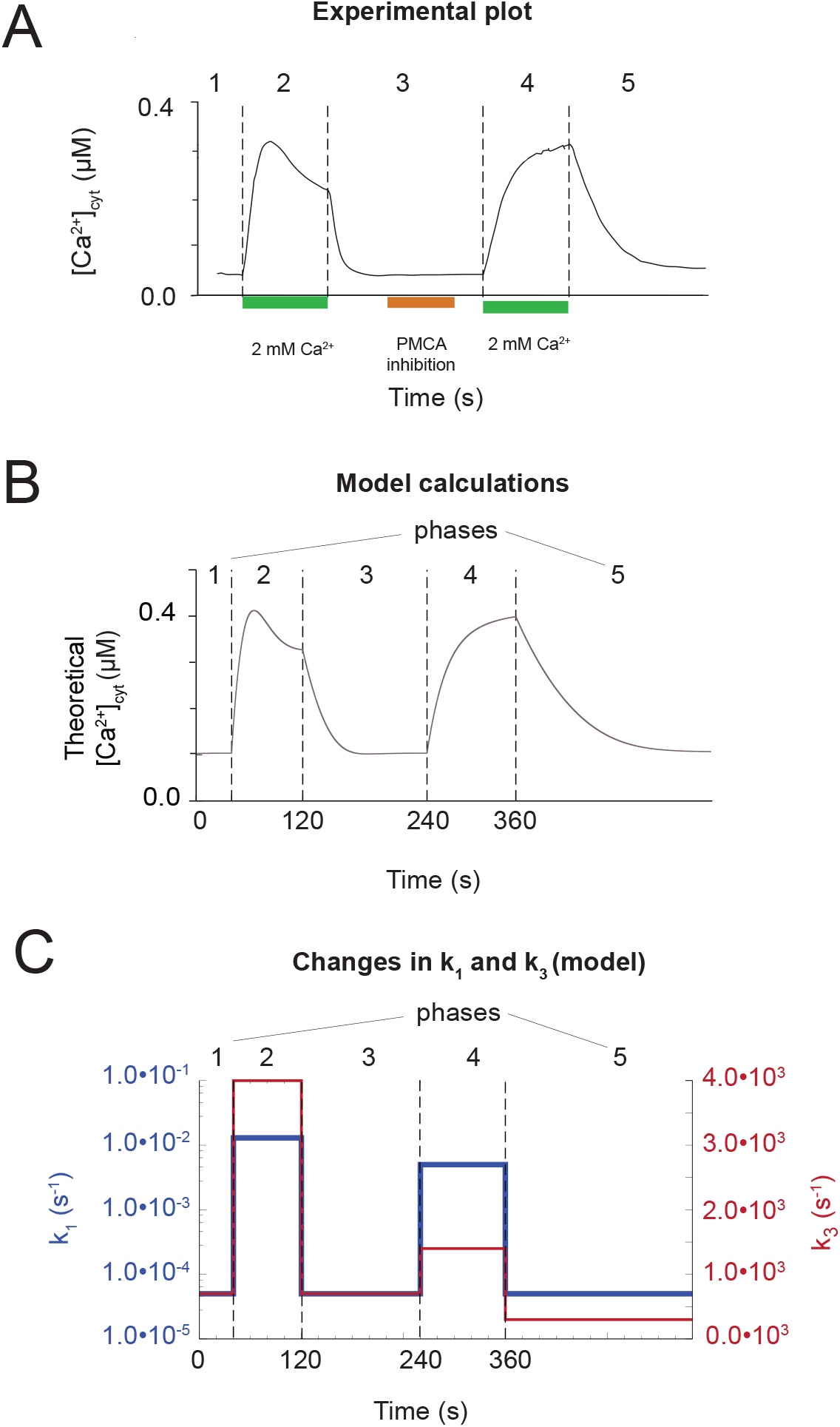
PMCA induced 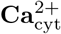 dynamics before and after its inhibition. Panel A shows a redrawn figure of Sedova and Blatter’s experimental results (Fig 1A in [59]) showing the dynamics of cytosolic Ca^2+^ upon capacitative Ca loading. The figure is divided into 5 phases indicated by dashed vertical lines. Phase 1: steady state of cytosolic Ca^2+^ prior to extracellular Ca treatment; phase 2: cells are treated with 2 mM extracellular Ca^2+^; phase 3: external Ca^2+^ is washed out and PMCA is inhibited by carboxyeosin; phase 4: 2 mM extracellular Ca^2+^ is re-applied; phase 5: extracellular Ca^2+^ is washed out. Panel B shows our model’s response in cytosolic calcium when the inflow of calcium into the cell (*k*_1_) and the activity of PMCA (*k*_3_) undergo step-wise changes. Panel C shows the applied changes in *k*_1_ and *k*_3_ during the five phases to mimic the experimental Ca profiles. Phase 1: *k*_1_=5 × 10^−5^s^−1^, *k*_3_=7.0 × 10^2^s^−1^; phase 2: *k*_1_=1.3 × 10^−2^s^−1^, *k*_3_=4.0 × 10^3^s^−1^; phase 3: *k*_1_=5 × 10^−5^s^−1^, *k*_3_=7.0 × 10^2^s^−1^; phase 4: *k*_1_=5 × 10^−5^s^−1^,*k*_3_=1.4 × 10^3^s^−1^; phase 5: 5 × 10^−5^s^−1^, *k*_3_=3.0 × 10^2^s^−1^. Other rate constant values, initial concentrations, and the model’s Fortran source file together with executables can be found in the Supporting Information ‘S3 Program’.

Another explanation could be that in absence of PMCA the Na^+^-Ca^2+^ exchanger NCX brings Ca^2+^ down to low concentrations similar to PMCA [59]. However, since NCX is considered to have a lower affinity for calcium and is generally believed to be active at higher Ca^2+^ concentrations in comparison to PMCA this suggestion remains controversial [15, 16].

### PMCA’s hysteretic behavior

However, the changes invoked on *k*_1_ and *k*_3_ alone are not sufficient to model the transients observed in the above experiments by Sedova and Blatter. There needs to be present an inherent slowness in PMCA’s response kinetics. In other words, PMCA acts as hysteretic enzyme, i.e. shows a slow activation/reactivation kinetics [59, 62, 63]. Frieden [63] studied this slowness of certain enzymes upon activations and termed the phenomena ‘hysteretic’ behavior. Scharff et al. [62] concluded that PMCA reacts hysteretically to increases in cytosolic Ca^2+^ inflow and that this is a necessary property to enable the occurrence of transients.

In general, transients are important in signaling and differences in the transient’s strength in amplitude or frequency (if oscillatory) are interpreted as different signals. For example, the slowness in PMCA activation allows the concentration of Ca^2+^ in the cytosol to increase to a relevant concentration for signaling purposes, while a slowness in its deactivation would give the PMCA time to function as an extrusion mechanism to bring the cytosolic Ca^2+^ level down to its set-point again.

As a way of incorporating hysteresis, we have employed the calmodulin (CaM) binding as the limiting factor, as it is widely agreed upon that CaM binding to PMCA isoforms can vary, and that CaM binding is notoriously slow. Those pump isoforms dominantly found in non-excitable cells are characterized as “slow PMCA pumps”, whereas in excitable cells where Ca^2+^ can vary frequently and rapidly, some isoforms of PMCA are characterized as “fast pumps” [64–66].

CaM is also described as a limiting factor in the cell for regulation of its targets. Some researchers report that most CaM in the cytosol exist as ApoCaM (unbound to Ca^2+^) at resting Ca^2+^ levels, and that in most cases ApoCaM does not bind to its targets as long as Ca^2+^ has not bound to it [67,68]. However, Wu and Bers note that some ApoCaM also binds to many proteins like ion channels, membrane proteins and receptors etc. either as part of their structure or for the purpose of local activation by Ca^2+^ binding. This means that upon Ca^2+^ binding, targets already bound to (Apo)CaM can immediately be activated, while the unbound targets need to compete for free Ca^2+^-CaM in the cytosol. This arrangement fits well with our model where ApoCaM (M in the model) first binds to PMCA (leading to PMCA•M in the model) before activation by Ca^2+^ occurs leading to the active pump form PMCA•M^*^. However, there are also arguments that PMCA competes for the free Ca^2+^-CaM complex [67, 69]. Nevertheless, it appears more likely that CaM binds to PMCA and Ca^2+^ as targets instead of being present as a free Ca^2+^-CaM complex in the cytosol: it has been estimated that only about 1 % of the total intracellular CaM exist as free Ca^2+^-CaM complexes [70]. In our model the portion of free Ca^2+^-CaM complex varies between about 4 – 10 % depending on the rate constants applied in the calculation, where most of them have been taken from literature. The total CaM concentration is set to 10 *μ*M in the model [67]. Rate constant *k*_9_ for the off-rate, i.e. the dissociation of M·Ca_4_ into M (CaM) and Ca^2+^, is 5.0 s^−1^ and *k*_8_ for the on-rate, for the association of Ca^2+^ and M (CaM) to M·Ca_4_, is set to 2.5 *μ*M^-1^ ·s^-1^ in order to keep the K_*d*_ at a defined value of 2 *μ*M [70, 71].

In addition to the PMCA-CaM binding, a change in *k*_3_ can also add to the hysteretic behavior of the pump as seen in Fig 6C. The hysteretic properties of the pump regarding *k*_3_ was investigated in more detail when the NCX pump was added to the model (see next section).

### Implementing NCX

Next we added the Na^+^-Ca^2+^ exchanger (NCX) to the model. NCX moves one Ca^2+^ ion out of the cell with the exchange of 3 Na^+^ ions. Using bovine endothelial cells Sedova and Blatter [59] investigated the role of NCX when PMCA was inhibited by La^3+^ ions, and when NCX was inactivated by Na^+^ free conditions. They found that under Na^+^ free conditions and in the presence of La^3+^, i.e. when both PMCA and NCX are not active, cytosolic Ca^2+^ cannot reach resting levels. However, when Na^+^ was added in presence of La^3+^, i.e. when only NCX was active, Ca^2+^ concentrations were able to reach resting levels, but more slowly. These results indicate that NCX alone is able to reduce cytosolic Ca^2+^ to resting levels in the absence of PMCA.

Based on the findings by Chou [72], we have included calmodulin (M in model) as a regulator of NCX activation by Ca^2+^. In this way, the NCX is modeled analogous to PMCA, i.e. NCX is complexed with calmodulin (NCX•M) with the activated form (NCX•M^*^). Fig 7A shows, outlined in red, the added NCX part to the model.

**Fig 7.**
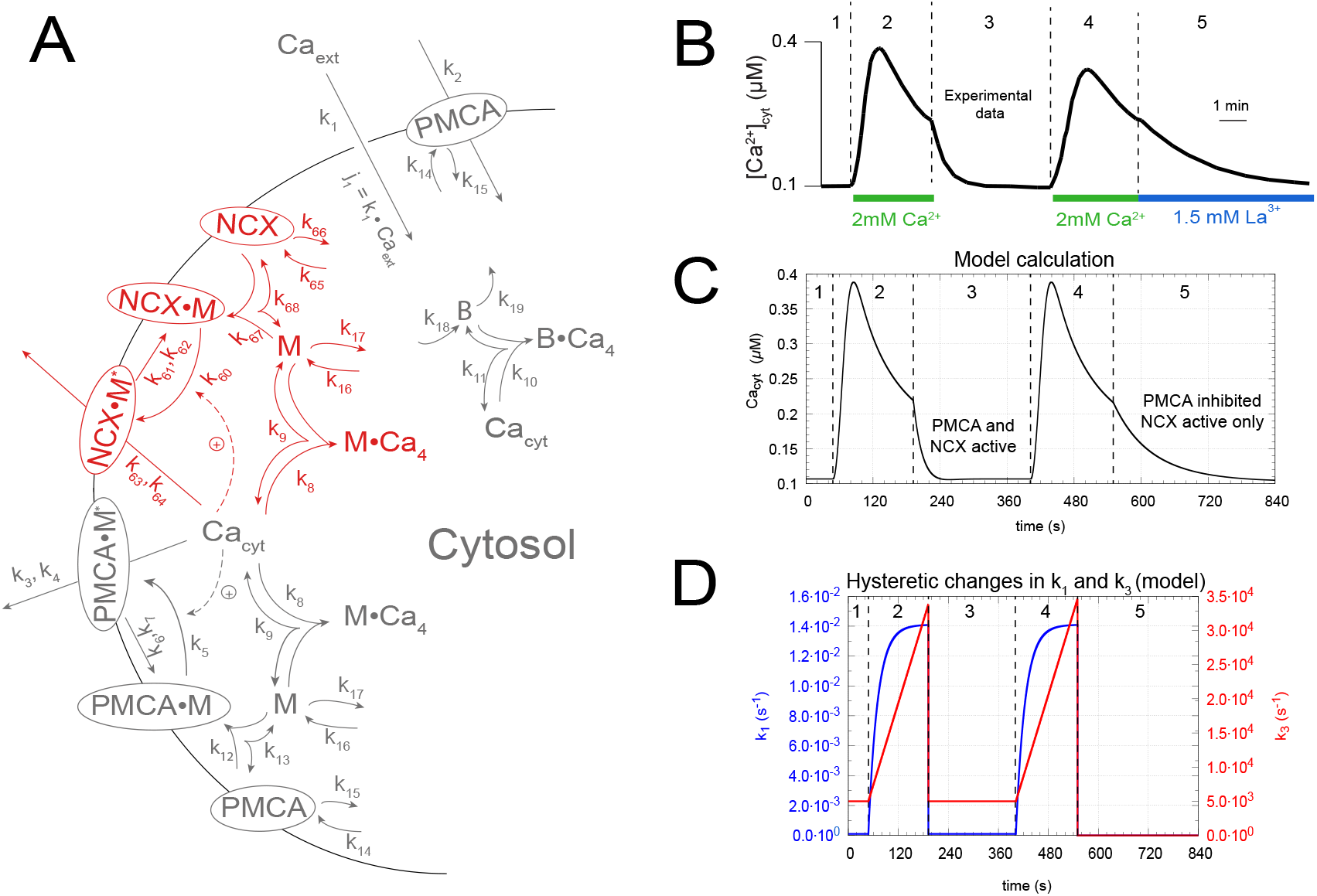
Cytosolic calcium concentration changes during PMCA-NCX based and NCX-only based calcium efflux. Panel A shows the model with NCX incorporated (outlined in red). The rest of the model is seen in grey representing the reactions that were already introduced. Panel B shows the experimental results redrawn from Sedova and Blatter (see Figure 6A in [59]). Panel C shows the corresponding model calculations. Phases 1 and 3: The cell is at resting state with both PMCA and NCX active. Phases 2 and 4: Cells are treated with 2 mM extracellular Ca^2+^ - in the model this change is represented by an increase of *k*_1_ (see text). Phase 5: Only NCX is active, PMCA is inhibited by La^3+^ - in the model this is done by setting *k*_3_ to zero. Panel D shows the (hysteretic) changes in the parameters *k*_1_ and *k*_3_ during phases 2 and 4 described by Eqs 10, 11, 12, and 13. Other rate constants, initial concentrations, and instructions how to run the model are found in Supporting Information ‘S4 Program’.

Thus, similar to Ca^2+^ regulation by PMCA, a set-point for cytosolic Ca^2+^ can also be formulated for NCX. Setting the rate equation of the calcium-activated calmodulin-associated form (*NCX*·*M* ^*^) to zero,

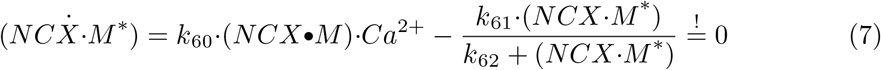

assuming that *k*_62_ ≪ (*NCX*·*M* ^*^), and then solving from resulting Eq 8 for 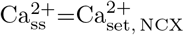

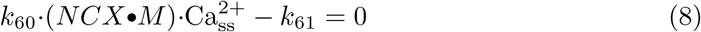

we get for the NCX-based set-point:

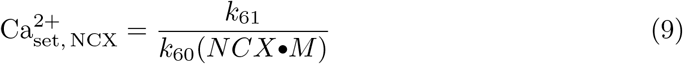

### Model calculations including NCX in comparison with experimental results

In comparison with the experimental results by Sedova and Blatter [59] (Fig 7B) we show here calculations with the inclusion of NCX. Since NCX is assumed to bind calcium less efficiently than PMCA the K_M_ of NCX was taken to be 100 *μ*M, which is approximately 2 orders of magnitude larger than the K_M_ with respect to calcium for PMCA. On the other hand, the turnover number of NCX (in the model described by *k*_63_) is set to 1 × 10^5^ s^−1^, which is two orders of magnitude larger than the turnover number (*k*_3_) of PMCA. Fig 7C shows the calculated cytosolic calcium concentration mimicking the experimental results by Sedova and Blatter. In the model we have, in addition, assumed time-dependent hysteretic behaviors both in the capacitative inflow of calcium into the cytosol (described by rate constant *k*_1_) and in the transport of Ca^2+^ by PMCA (described by rate constant *k*_3_). In phases 2 and 4 *k*_1_ is assumed to increase and reaching a maximum value according to the relationships:

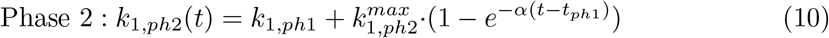

and

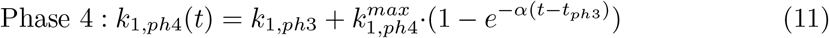

In Eqs 10 and 11 *k*_1,*ph*1_ and *k*_1,*ph*3_ are constants both having a value of 1×10^−4^s^−1^. 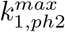 and 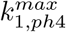 are the maximum values of *k*_1_ for respective phases 2 and 4 with a value of 1.4×10^−2^s^−1^ each. The parameter *α* is 4.5×10^−2^s^−1^. *t*_*ph*1_ and *t*_*ph*3_ denote the times when phases 1 and 3 end, respectively. When these *k*_1_-time dependencies are implemented into the model we get the *k*_1_ profile as shown in Fig 7D (outlined in blue). The hysteretic changes of *k*_3_ in phases 2 and 4 is described by the linear relationships

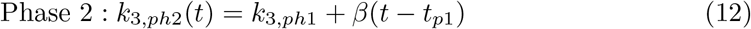

and

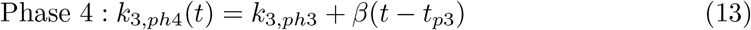

with *β*=200s^−2^ and *k*_3,*ph*1_=*k*_3,*ph*3_=5 × 10^3^s^−1^. In phase 5 *k*_1_ and *k*_3_ are zero, except *k*_63_ (turnover number of NCX), which for all five phases is 1 × 10^5^s^−1^. For a description of the other rate constants, initial concentrations, and how to run the program see Supporting Information ‘S4 Program’.

Comparing Figs 7B and C shows that the model calculations represent a good fit to the experiment. The model also shows that NCX can drive cytosolic Ca^2+^ concentrations down to resting values without PMCA present, but at a slower rate, just as observed in the experiments. This is mainly because of the difference between the *K*_*M*_ values of PMCA and the NCX making PMCA a high affinity pump with low *K*_*M*_ (=*k*_4_). The relative high *K*_*M*_ (=*k*_64_) of NCX makes NCX a low affinity pump. In order to get the peak shape of the cytosolic Ca^2+^ concentration during phases 2 and 4, the above hysteretic changes of Ca^2+^ inflow (via SOCC, represented here by *k*_1_) and Ca^2+^ outflow (via PMCA *k*_3_) appear necessary. Possibly, also NCX may exhibit hysteretic behavior, but this was not included.

In an additional set of experiments (Fig 8A) Sedova and Blatter investigated the sequential and simultaneous inhibition of PMCA and NCX. While NCX was inhibited using Na^+^-free conditions cells were treated with 2mM extracellular Ca^2+^. The increase and decrease of cytosolic calcium indicate an active PMCA pump (phase 2, Fig 8A). Then, in phase 3 extracellular Ca^2+^ was washed out and cells were treated with La^3+^ to inhibit PMCA while still having Na^+^-free conditions (Fig 8A). Since both PMCA and NCX were now inactive cytosolic calcium levels did not change much (phase 3, Fig 8A). Finally, in phase 4 cells were treated with sodium ions, which activated NCX and drives cytosolic calcium levels down to resting values close to 0.1*μ*M. This clearly indicates that NCX was responsible for the decrease of cytosolic calcium in phase 4.

**Fig 8.**
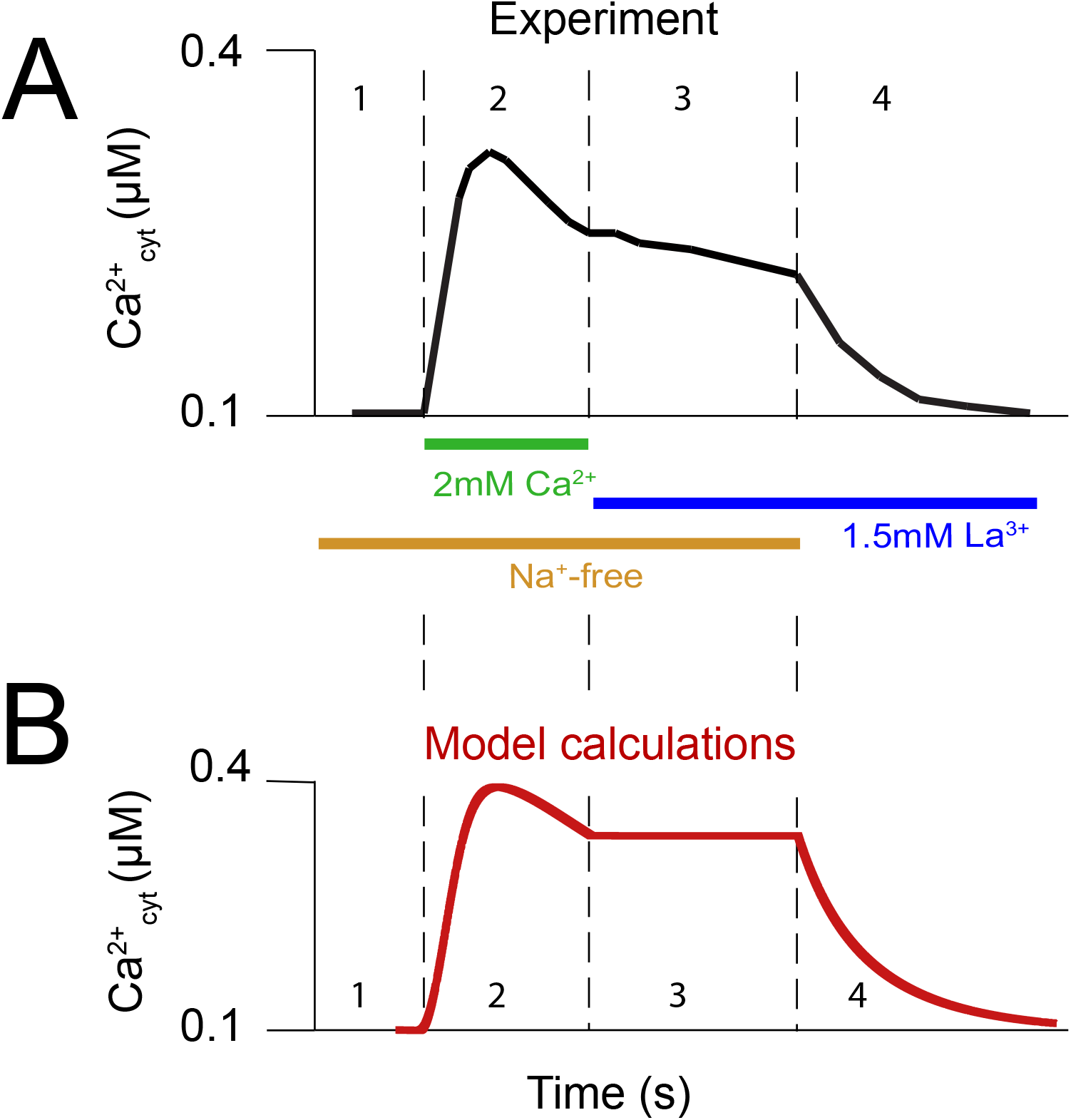
Ca^2+^ efflux following La^3+^ inhibition of PMCA. Panel A is redrawn from Sedova and Blatter’s experimental record (Fig 6B in [59]). Panel B derives from a model calculation (S5 Program) simulating the experimental results in A. Phase 1: Cells are at resting conditions, no Ca^2+^ enters the cytosol. Only PMCA is active, NCX is inactive due to Na^+^-free conditions. In the model inactive NCX is achieved by setting *k*_63_ to zero. Phase 2: Cells are treated with 2mM extracellular calcium and the capacitative inflow of calcium into the cytosol starts. In the model *k*_1_ and *k*_3_ increase hysteretically (Eqs 10 and 12, S5 Program). Only PMCA is considered to be responsible for pumping calcium out of the cytosol. Phase 3: Extracellular calcium is removed and cells are treated with La^3+^ inhibiting PMCA. Since the medium is still sodium-free NCX is inactive. In the model both *k*_3_ and *k*_63_ are set to zero. Phase 4: NCX is activated by adding a sodium salt solution to the medium, while PMCA is still inactive due to the presence of La^3+^. In the model NCX activation is achieved by setting *k*_63_=1 × 10^5^s^−1^. Initial concentrations, other rate constant values, and how to run the model are described in Supporting Information ‘S5 Program’.

Fig 8B shows a corresponding calculation of the model mimicking the results by Sedova and Blatter. The experiments show that the La^3+^-treated cells in phase 3 still show a slight decrease in cytosolic calcium, indicating that the the PMCA inhibition was not 100% perfect or that there may be a still unknown constitutive calcium outflow from the cytosol.

### Role of ER in calcium regulation

#### Overview

The inflow and outflow of Ca^2+^ between the endoplasmatic reticulum (ER) and the cytosol plays a major role in the calcium homeostasis in the ER as well as in the cytosol. The ER functions as a calcium store with concentrations comparable to extracellular calcium. Furthermore, the ER is heavily involved in calcium-mediated signaling (Ca^2+^ inflow into the cytosol) through the inositol 1,4,5-triphosphate receptor IP_3_R and ryanodine receptor RyR located in the ER membrane (Fig 1). The activity of the IP_3_-bound IP_3_R channel (IP_3_R•IP_3_, Fig 3) shows a bell-shaped activity profile with a maximum at around pCa 7 [73]. The IP_3_R•IP_3_ channel is activated by low cytosolic Ca^2+^ concentrations, i.e. when pCa *>* 7, while inhibition occurs at higher cytosolic concentrations, i.e. when pCa *<* 7. An analysis of the experimental data by Kaftan et al. [73] (shown below) indicates an asymmetric channel activity profile with a stronger Ca^2+^-inhibition cooperativity around 2, while the activation cooperativity is found to be approximately 1.

When the calcium content in the ER is low stromal interaction molecules 1 and 2 (STIM1 and STIM2, represented as STIM in the model, Fig 3) activate store operated Ca^2+^ channels (SOCCs), which lead to an inflow of Ca^2+^ into the cytosol. Subsequently, the ER is refilled through the sarco-endoplasmatic reticulum Ca^2+^ ATPase (SERCA), an ATPase similar to PMCA.

There is evidence that ER-luminal Ca^2+^ leaks into the cytosol. Analyzing the data by Luik et al. [74] shows a good agreement with a first-order kinetic Ca depletion of the ER with a rate constant of 0.012 s^−1^ and an initial leakage velocity of 0.36 *μ*M/s (S6 Program). The leak data recorded by Camello et al. [10] shows almost zero-order Michaelis-Menten kinetics with a slight hysteretic increase of V_max_ and an average leakage rate of 0.22 *μ*M/s (S6 Program).

The ER also contains several high-capacity Ca^2+^-binding proteins, such a calreticulin [75, 76], which represent ER’s (and other organelle’s) overall Ca^2+^ storage capacity. According to Michalak et al. (see [75], and references therein) one calreticulin molecule can bind between 20-50 Ca^2+^ ions. In the model (Fig 3) we have represented these high-capacity storage proteins inside the ER with the letter L and enabled them to bind 30 Ca^2+^ ions.

#### Store operated (capacitative) Ca^2+^ entry

Fig 9 shows, outlined in red, the reaction scheme of the capacitative Ca^2+^ entry in the model determined by the Ca^2+^ concentration in the ER (Ca_lum_). For simplicity, only one STIM form has been considered in the model. Biologically, there have been found two STIM forms (STIM1 and STIM2), and even though STIM1 and STIM2 have a similar binding affinity towards Ca^2+^ (STIM1: K_*d*_ 0.2-0.6 mM, STIM2: K_*d*_ 0.5 mM) they display differences in the regulation of SOCC possibly due to structural reasons [77, 78]. STIM2 responds to small changes of Ca^2+^ [77]. The results from Zheng et al. [77] indicate that STIM1 is the only Ca^2+^ sensor responsible for SOCC activation, while STIM2 seems only to play a minor role. On the other hand, Brandman et al. [12] investigated the effects of smaller decreases in luminal Ca^2+^ concentration and suggested that also STIM2 takes part in the feedback regulation of calcium in the cytosol and ER. These findings are supported by the work of Thiel et al. [79]. In the model, when STIM dissociates at low Ca_lum_ concentrations from the STIM•Ca complex (Fig 9), STIM concentration increases and activates the Ca^2+^ inflow through nearby SOCC channels [74, 80].

**Fig 9.**
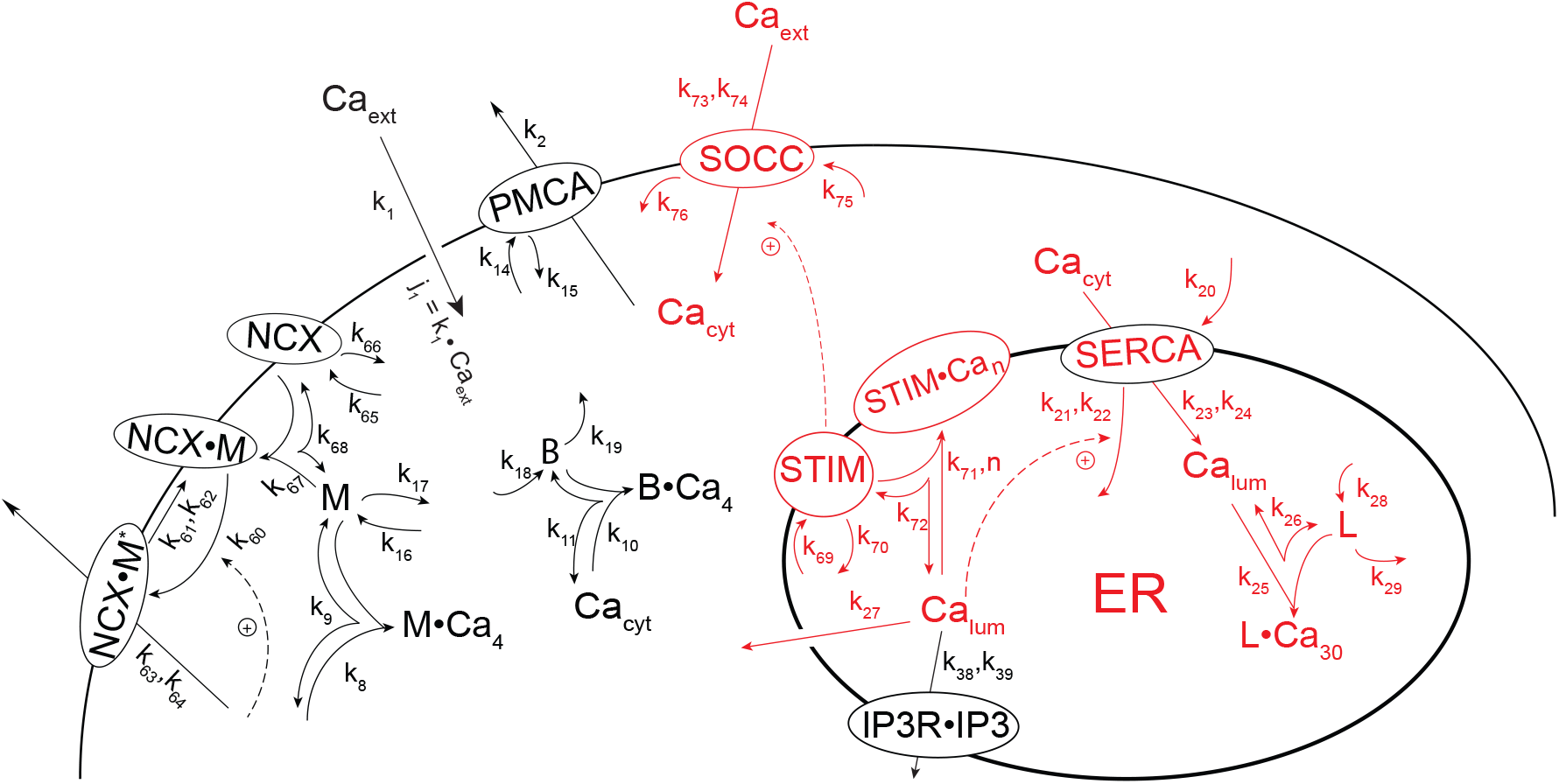
Capacitative Ca^2+^ entry. Outlined in red is the scheme used in the model. Decreasing Ca concentrations in the ER (Ca_lum_) lead to a dissociation of the STIM•Ca complex and to increased STIM levels activating nearby SOCC channels. The parameter *n* describes the Ca_lum_-cooperativity when binding to STIM. To see the relationship between *j*_*SOCC*_ at different Ca_lum_ concentrations by STIM activation, the SERCA channel has been formulated as an inflow controller [28] for Ca_lum_, which allows to keep Ca_lum_ constant at different levels by changing *k*_20_. For details, see text and S7 Program.

Luik et al. [74] studied the inflow rate of calcium into the cytosol by the SOCC channel (rate: *j*_*SOCC*_) as a function of the calcium concentration in the ER (Ca_lum_). We have analyzed the Luik et al. data (Fig 10) in terms of a Ca_lum_ derepression relationship, which activates *j*_*SOCC*_ via STIM (Fig 9), i.e., using the relationship

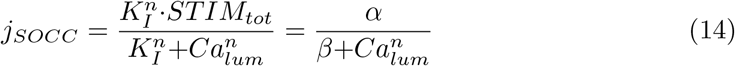

where *K*_*I*_ is an inhibition constant, *STIM*_*tot*_ is the total (constant) STIM concentration and *n* is the cooperativity (Hill-coefficient) of the *j*_*SOCC*_ inhibition by Ca_lum_. *α, β*, and *n* in Eq 14 were fitted to the experimental data by Luik et al. (S7 Program). Although Eq 14 is slightly different from the Hill-equation used by Luik et al. the obtained fit and Hill-coefficient *n* are comparable. While Luik et al. interpreted their Hill-coefficient of 4.2 as a sign of STIM1 oligomerization, we feel that in our case a Hill-coefficient of 3.59 appear misleading, although representing a “best-fit”. Our high Hill-coefficient may be misleading by two reasons: firstly, Eq 14 implies a derepression mechanism, where Ca_lum_ apparently inhibits *j*_*SOCC*_, but this is mechanistically only done indirectly via binding to STIM. Secondly, the fit in Fig 10A (green line) indicates that the slope d*j*_*SOCC*_/d(Ca_lum_) should be reaching a plateau at low Ca_lum_ concentrations, which is not strongly supported by the data. We have therefore analyzed the Luik et al. data in terms of a cooperative reversible Ca_lum_ binding to STIM with Hill-coefficient *n*, described by the process (see S1 Text for the rate equations):

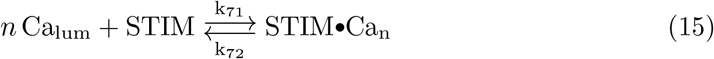

**Fig 10.**
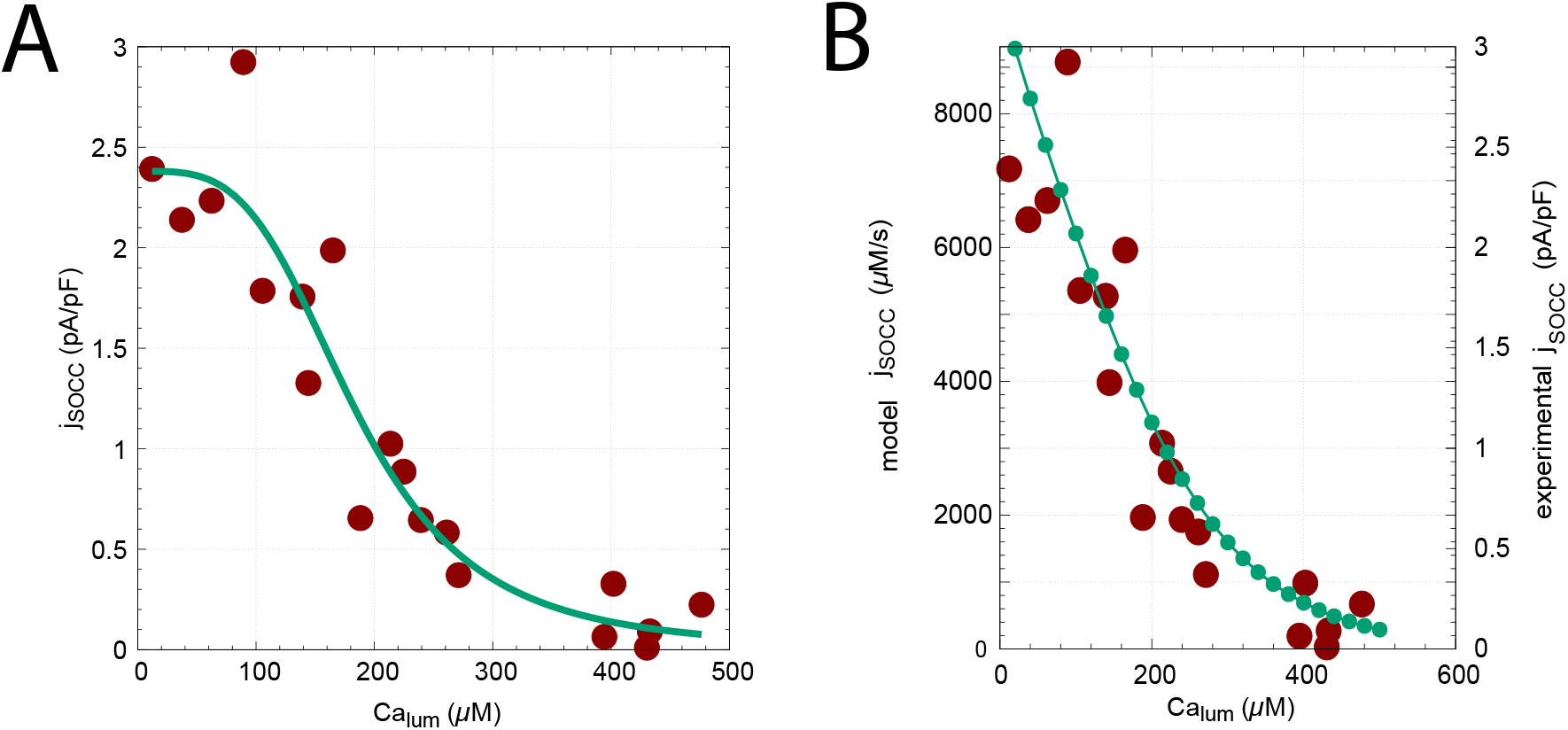
Capacitative Ca^2+^ inflow rate *j*_*SOCC*_ as a function of calcium concentration in the ER (Ca_lum_). Panel A: experimental results (red solid dots) redrawn after Fig 1c from Luik et al. [74]. Green line shows a nonlinear fit for a constant Hill-coefficient with *n*=3.59±0.9 (Eq 14). For details, see S7 Program. Panel B: model calculation using Eq 14 with a variable Ca_lum_ cooperativty *n* changing linearly from n=0.8 at Ca_lum_=20 *μ*M to *n*=1.3 at Ca_lum_=500*μ*M. For details, see S7 Program.

By successively changing Ca_lum_ from 20*μ*M to 500*μ*M (by steps of 20*μ*M) we found that an eye-balled “best fit” to the Luik et al. data can be obtained (Fig 10B) when *n* is changed linearly from 0.8 at Ca_lum_=20*μ*M to *n*=1.3 at high Ca_lum_=500*μ*M. We interpret the changing Hill-coefficent as a hysteretic/conformational effect. In other words, at low Ca_lum_ concentrations not all Ca-binding sites are available for binding to Ca, while at higher Ca_lum_ concentrations conformational changes may expose more Ca-binding sites such that more than one Ca^2+^ ion can bind to STIM. Such a view is in agreement with the review by Grabmayr et al. [81], which indicates that some STIM isoforms can bind more than one Ca^2+^ ion.

In the model the steady state of Ca_lum_ is determined by the SERCA channel, which has been ‘wired’ in form of a negative feedback [28] inflow controller. The amount of SERCA is increased by *k*_20_, while Ca_lum_ signals a zero-order decrease of SERCA, i.e.

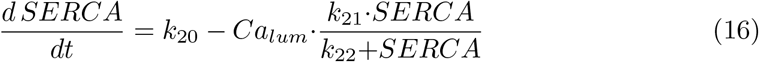

At steady-state (i.e., 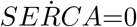) and zero-order condition with respect to SERCA, (*k*_22_≪(*SERCA*)), the *Ca*_*lum*_ set-point is determined by Eq 16:

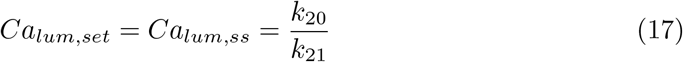

Fig 11 illustrates the robustness of the Ca_lum_ set-point by calculating *Ca*_*lum,ss*_ at five different phases (1-5) in which *k*_20_ is changed from *k*_20_=1000*μ*M/s (phase 1) to 200*μ*M/s (phase 5) by steps of 200*μ*M/s. Panel A shows the decrease in Ca_lum_ concentration when *k*_20_ is successively decreased, while panel B shows the corresponding increase of *j*_*SOCC*_. See S7 Program for details.

**Fig 11.**
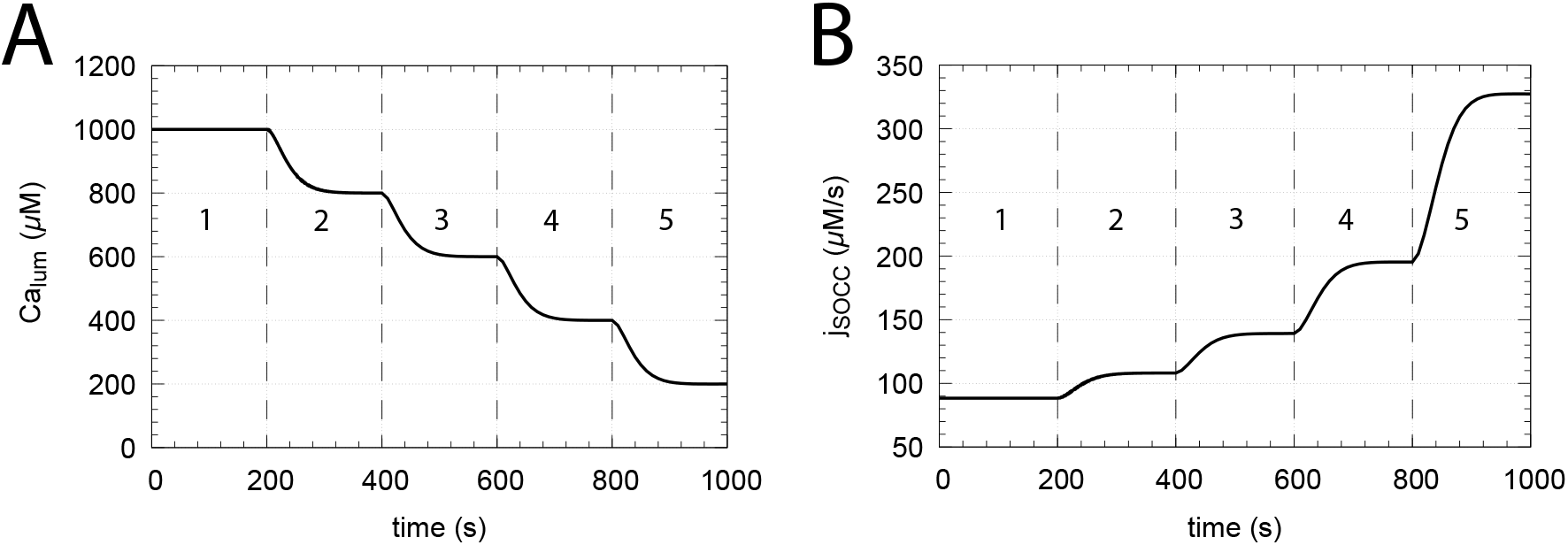
SERCA inflow control determines Ca_lum_ concentration. The figure illustrates the decrease in *Ca*_*lum*_ concentration as its set-point (Eq 17) decreases with changing *k*_20_ together with the corresponding up-regulation of *j*_*SOCC*_ as *Ca*_*lum*_ gets lower. Panel A: Five phases 1-5 show the changed Ca_lum_ concentrations when *k*_20_ values are successively decreased. Phase 1: *k*_20_=1000*μ*M/s, phase 2: *k*_20_=800*μ*M/s, phase 3: *k*_20_=600*μ*M/s, phase 4: *k*_20_=400*μ*M/s, and phase 5: *k*_20_=200*μ*M/s. *k*_21_=1s^−1^ in all five phases. Panel B: corresponding increase of *j*_*SOCC*_ in response to the Ca_lum_ concentration changes in panel A. For details, see Supporting Information S7 Program.

We have revisited the experiments by Sedova and Blatter (Ref [59] and Fig 7) and calculated Ca_cyt_ concentrations by using the capacitative Ca inflow mechanism from Fig 9. Fig 12 shows the obtained Ca_cyt_ concentrations (panel A, phases 2 and 4) when cells are exposed to 2mM external Ca at zero *Ca*_*lum*_ levels. During phase 5 PMCA is inhibited and only NCX is active.

**Fig 12.**
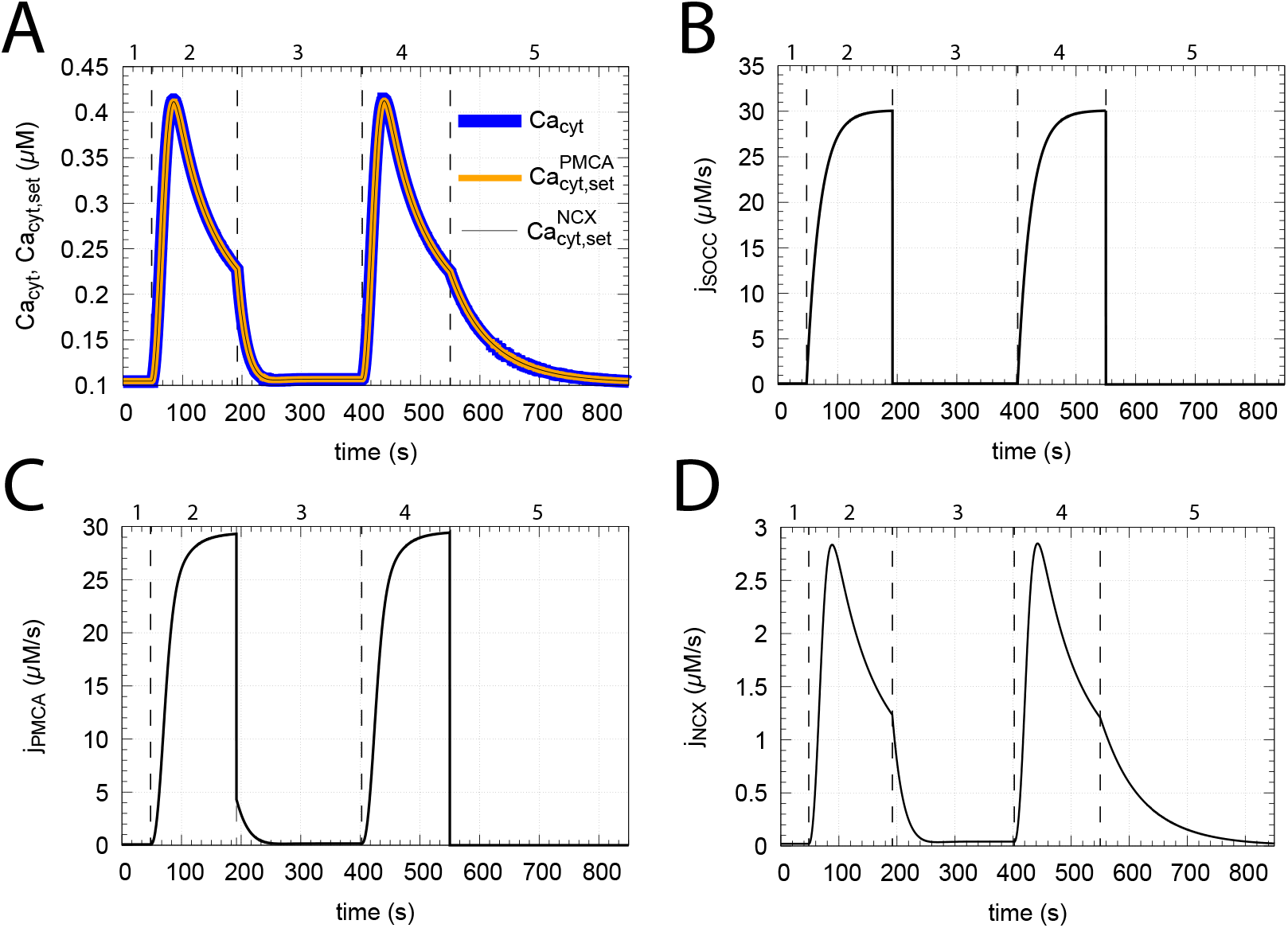
Cytosolic Ca^2+^ changes due to capacitative Ca inflow. Computations were performed with the mechanism outlined in Fig 9. In phases 2 and 4 extracellular Ca was set to 2 mM along with no Ca in the ER (*Ca*_*lum*_=0*μ*M) and a fully-inhibited SERCA (*k*_23_=0s^−1^). Panel A: calculated Ca_cyt_ and set-points for PMCA and NCX as a function of time. Panels B and C: SOCC and PMCA are imposed to undergo hysteretic changes in phases 2 and 4, such that it takes some time until maximum channel activities are established. Panel D: the channel turnover number of NCX *k*_63_ is kept high (1 × 10^5^s^−1^) but constant for all five phases. For details, see Supporting Information S7 Program.

To achieve the Ca_cyt_ profiles as in Fig 7 we assumed that SOCC and PMCA show hysteretic behaviors, i.e., for both channels it takes some time until their maximum activities are established. No hysteretic effects are imposed on NCX. Here, the channel turnover number is kept high and constant at *k*_63_=1×10^5^s^−1^ throughout all five phases.

#### Ca_cyt_-induced calcium release through IP_3_R and its inhibition at high Ca_cyt_

We have chosen to include IP_3_R as the only Ca^2+^ release channel in the ER membrane. This is because the RyRs are located in the sarcoplasmic reticulum of excitable cells and are involved in events like the excitation-contraction coupling in both cardiac and skeletal muscle [82]. Also of relevance is the fact that IP_3_R alone has been found to produce agonist-induced Ca^2+^ oscillations. A bit contradictory, Sanders showed in 2001 that both IP_3_R and RyR were needed for sufficient oscillatory behavior in vascular smooth muscle cells (VSMCs) [83]. However, two years later, in 2003, McCarron found that even if RyRs are blocked IP_3_R alone can sufficiently produce agonist-induced Ca^2+^ oscillations [84]. These results were from excitable cells, which could also justify the usage of only the IP_3_R channel to move Ca from a store (here ER) into the cytosol. In this work we have not primarily been focussed on oscillations, but at the end of the paper we show that the concerted action of the SOCC-SERCA-IP_3_R axis together with PMCA/NCX can result in sustained oscillations.

IP_3_R is regulated by both IP_3_ and Ca_cyt_, where the regulation of IP_3_R by Ca_cyt_ is biphasic and results in a bell-shaped IP_3_R activity profile with a maximum at about 100-300 nM Ca_cyt_ (pCa_cyt_≈6.5-7) [73, 85–88]. The Ca^2+^ release from the store (ER) due to Ca_cyt_ activation is termed Calcium-Induced Calcium-Release (CICR) [5, 89, 90]. IP_3_ originates from an extracellular signal (e.g. hormones, growth factors, light, odorants, neurotransmitters) stimulating a G protein-coupled receptor in the cell membrane, which activates phospholipase C (PLC), and hydrolyzes phosphatidylinositol 4,5-bisphosphate (PIP2) into IP_3_ and diacylglycerol (DAG) [89–92]. The IP_3_R channel has three isoforms, but in this model we have chosen to have only one type of IP_3_R. Physiologically, IP_3_R1 is the most common and widespread form throughout cell types. All IP_3_R’s have a high affinity binding to IP_3_ with a *K*_*d*_ varying from about 14 to 163 nM. Some variations are found, but type 1 has been found to have a *K*_*d*_ between 50 - 89 nM [93–96]. All IP_3_R isoform activities display a bell-shaped Ca_cyt_ dependency. It is type 1 which has its peak at around 300 nM Ca_cyt_. We use this peak value in our model in agreement with experimental studies [73, 85–88, 94, 97].

IP_3_R’s large size made it difficult to study its structure, but advances in this direction have been made [22, 92, 98]. The structure based on the work by Fan et al. [22] is presented in Fig 13 together with the locations of the binding sites for IP_3_ and Ca^2+^ as suggested by Sienaert et al. and Ding et al. [99, 100].

**Fig 13.**
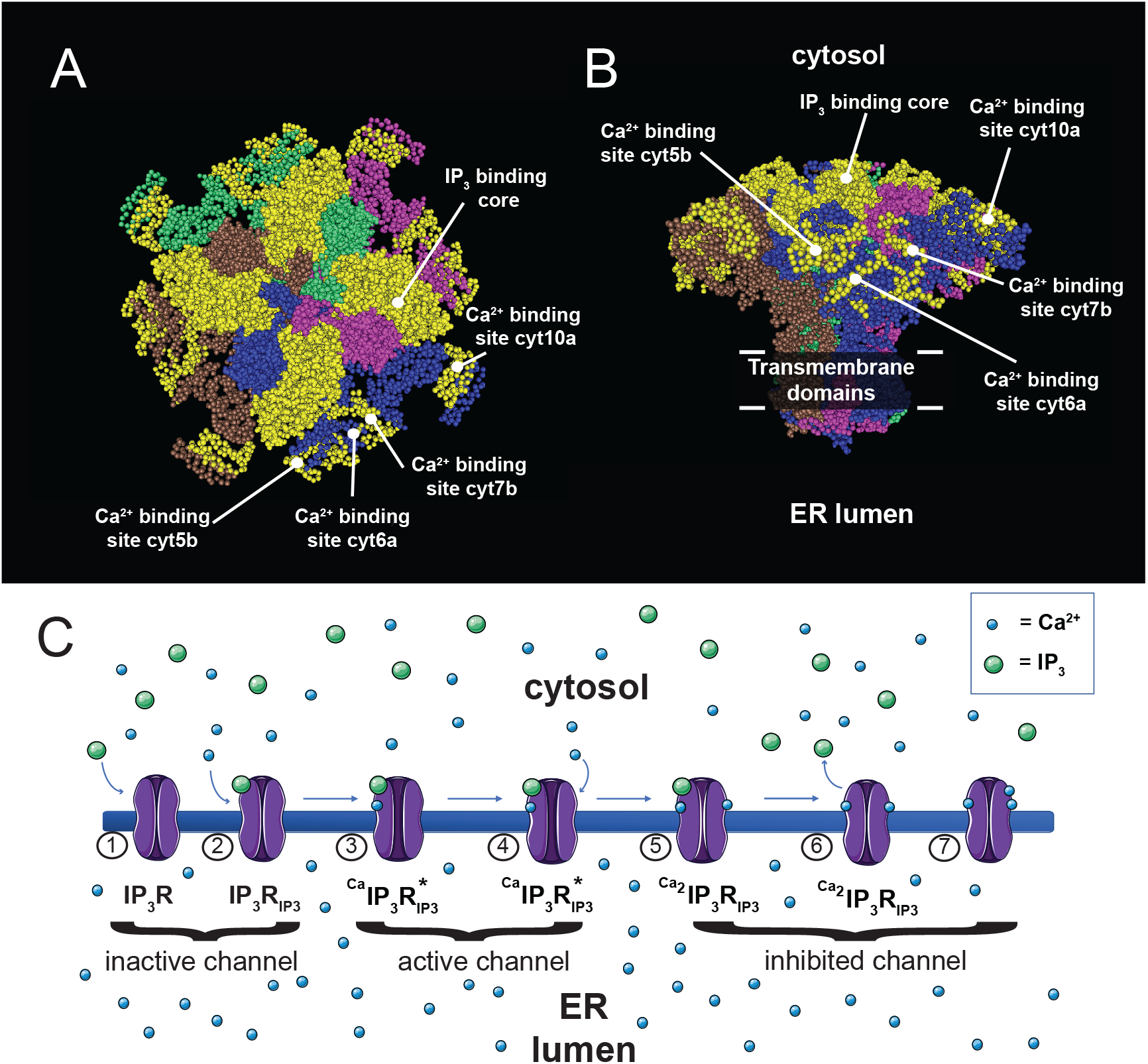
IP_3_R structure and mechanism of its activation and inhibition. Panels A and B show the IP_3_R channel structure based on the Protein Data Bank entry 3JAV (www.rcsb.org) from rat [22]. The four isomers are shown in blue, brown, purple and green color. Amino acid (aa) sequences of the Ca^2+^ binding sites and the IP_3_ binding core (IBC) were taken from Sienaert et al. and Ding et al., respectively, and are highlighted in yellow [99, 100]. Panel A shows the top view, while panel B gives a side view of the channel. Panel C shows an illustration of how IP_3_R channel may be activated and inhibited in a Ca^2+^ and IP_3_ dependent manner. Numbers 1-4 indicate the transition from an inactive to an active channel when Ca^2+^ and IP_3_ bind. Numbers 5-7 indicate the mechanisms of channel inhibition by the additional binding of Ca^2+^ and/or the dissociation of IP_3_ from IP_3_R•IP_3_.

IP_3_R gating, i.e. its activation by IP_3_ and Ca^2+^ as well as its inhibition by Ca^2+^ has been vastly investigated. Nevertheless, the mechanisms behind Ca-induced pore opening and closure are still not well understood and are somewhat controversial [89, 92, 101]. IP_3_R activation and inhibition by Ca^2+^ appear to be the result of conformational changes once Ca^2+^ and IP_3_ bind. There are currently 8 suggested Ca^2+^ binding sites on IP_3_R which are distributed over different structural domains. Their roles with respect to activating or inhibiting IP_3_R is not known [88, 89, 92, 101]. A calcium sensor region (Cas) has been suggested to be responsible for channel modulation by Ca^2+^ [98]. Concerning the inhibition of IP_3_R by Ca^2+^ there has also been the question whether the inhibition is due to a direct Ca^2+^ binding to an inhibitory site or if inhibition happens in association with a protein like calmodulin. On the other hand, studies have shown that calmodulin is not needed for Ca^2+^ inhibition, even though the Ca^2+^-calmodulin complex can inhibit the channel. Mutant studies lacking the high affinity Ca^2+^-calmodulin binding site displays a bell shaped Ca^2+^ dependence similar to the wild type. This indicates that Ca^2+^ itself or by the help of a different protein than calmodulin can inhibit IP_3_R [102, 103]. In a proposed model how Ca^2+^ influences IP_3_R Taylor and Tovey [89] and Taylor and Prole [104] suggest that IP_3_ makes the activating or inhibiting binding site accessible for Ca^2+^ dependent where IP_3_ binds on IP_3_R. Fig 13C gives an overview of the presumed activation and inhibition steps of IP_3_. It was also proposed that luminal Ca^2+^ might be tuning the channel in a way that it is less sensitive to IP_3_ when ER is depleted for calcium [89, 105, 106].

Based on the scheme in Fig 13C we have constructed a simple model (“dicalcic model”, Fig 14) which can describe the bell-shaped form of IP_3_R activity in response to the cytosolic calcium concentration Ca_cyt_.

**Fig 14.**
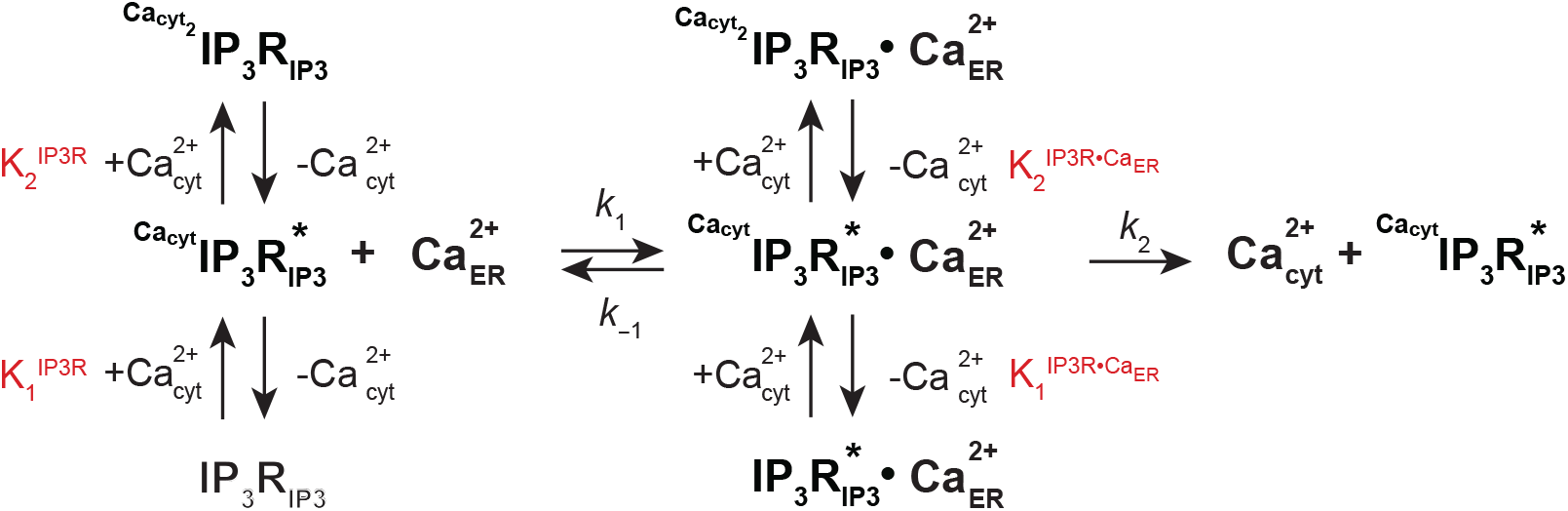
Dicalcic model. In this model the IP_3_R•IP_3_ channel has two cytosolic Ca binding sites, one that activates IP_3_R•IP_3_ at low cytosolic Ca^2+^ concentrations, while a binding at high cytosolic Ca^2+^ concentrations to the second site inhibits IP_3_R•IP_3_. The active form of the channel which transports Ca out of the ER into the cytosol is indicated by an asterisk. The four dissociation constants are outlined in red.

The model is analogous to the so-called diprotic model, which describes bell-shaped activity profiles of enzymes in dependence to pH [107]. In the dicalcic model we assume that IP_3_ is always bound to IP_3_R. We further assume that we have three IP_3_R forms: At low Ca_cyt_ concentrations (pCa_cyt_≫7) no Ca_cyt_ is bound to IP_3_R•IP_3_ and the channel is inactive (closed). At intermediate Ca_cyt_ concentrations (pCa_cyt_≈7) Ca_cyt_ binds and activates IP_3_R•IP_3_. This form is now able to transport calcium out of the ER and into the cytosol. At high Ca_cyt_ concentrations (pCa_cyt_≪7) there is further binding of Ca_cyt_ to IP_3_R•IP_3_ which leads to its inhibition.

We further assume that the binding between Ca_cyt_ and the IP_3_R channel can be formulated by four rapid equilibria

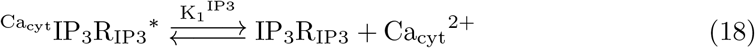

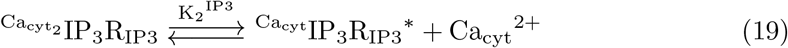

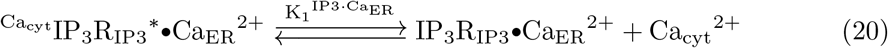

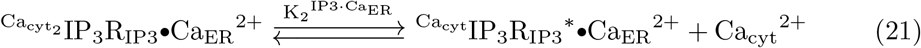

with dissociation constants 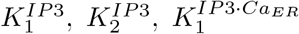, and 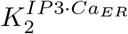. The asterisk in 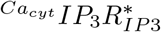 denotes the active form of the transporter, where one Ca_cyt_ has bound to IP_3_R (indicated by the left superscript ‘*Ca*_*cyt*_’). The right subscript ‘IP3’ in 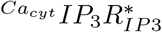 indicates the bound IP_3_. The active form 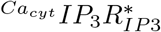 can bind Ca_ER_ <u>and</u> transport it into the cytosol (with channel turnover number *k*_2_, Fig 14). Although 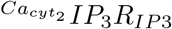 can bind Ca_ER_ and lead to 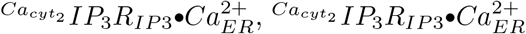 is not an active transport form (see Fig 14). The transport rate *v*_*IP*3*R*_ of the IP_3_R•IP_3_ channel is expressed as:

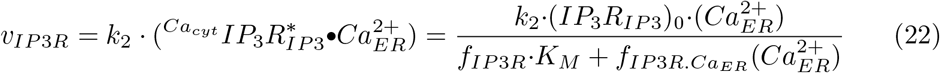

(*IP*_3_*R*_*IP*3_)_0_ is the total concentration of the IP_3_R•IP_3_ channel, which, for the sake of simplicity is considered to be constant. The factors *f*_*IP*3*R*_ and 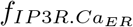 are analogous to the Michaelis acidity functions [107] and defined as:

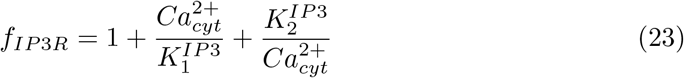

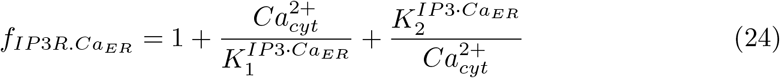

The *K*_*M*_ in Eq 22 is given as:

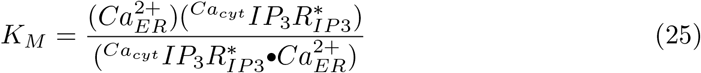

As anticipated, in Eq 22 cytosolic calcium acts both as an inhibitor and activator. The inhibition and activation terms are given by, respectively, 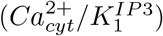 and 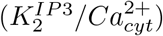 in Eq 23, while they are in Eq 24 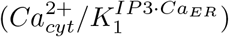 and 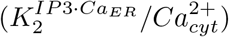. Thus, the *K*_1_’s take the form of inhibition constants, while the *K*_2_’s are activation constants.

We have used the experimental data by Kaftan et al. [73] to estimate the parameters in Eq 22. Kaftan et al. studied the Channel Open Probability of IP_3_R•IP_3_ as a function of Ca concentration at four different IP_3_ levels. We extracted the data (Fig 2 in [73]; see also S3 Text) and recalculated the experimental data as a function of Channel Open Probability versus pCa. Fig 15 shows the experimental data together with two fits at 2 *μ*M IP_3_. Panel A shows a fit of Eq 22 to the Kaftan et al. data when *f*_*IP*3*R*_ and 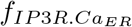 are taken from Eqs 23 and 24, respectively. We see that the experimental data (outlined in red) decrease much more abrupt with increasing Ca_cyt_ concentrations (decreasing pCa_cyt_) than the model (outlined in blue). A significantly better fit is obtained in panel B (outlined in green) when the cooperativity (Hill-coefficient) *n* for the increasing 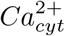 concentration in *f*_*IP*3*R*_ and 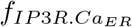 is increased to 2. In this case *f*_*IP*3*R*_ and 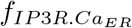 take the form:

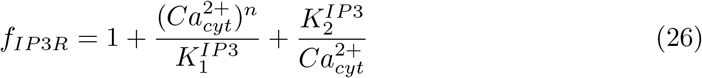

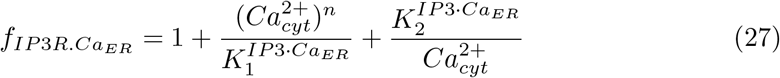

with *n*=2. Supporting Information ‘S3 Text’ shows the details of fitting the dicalcic model to all four IP_3_ concentrations used by Kaftan et al. [73].

**Fig 15.**
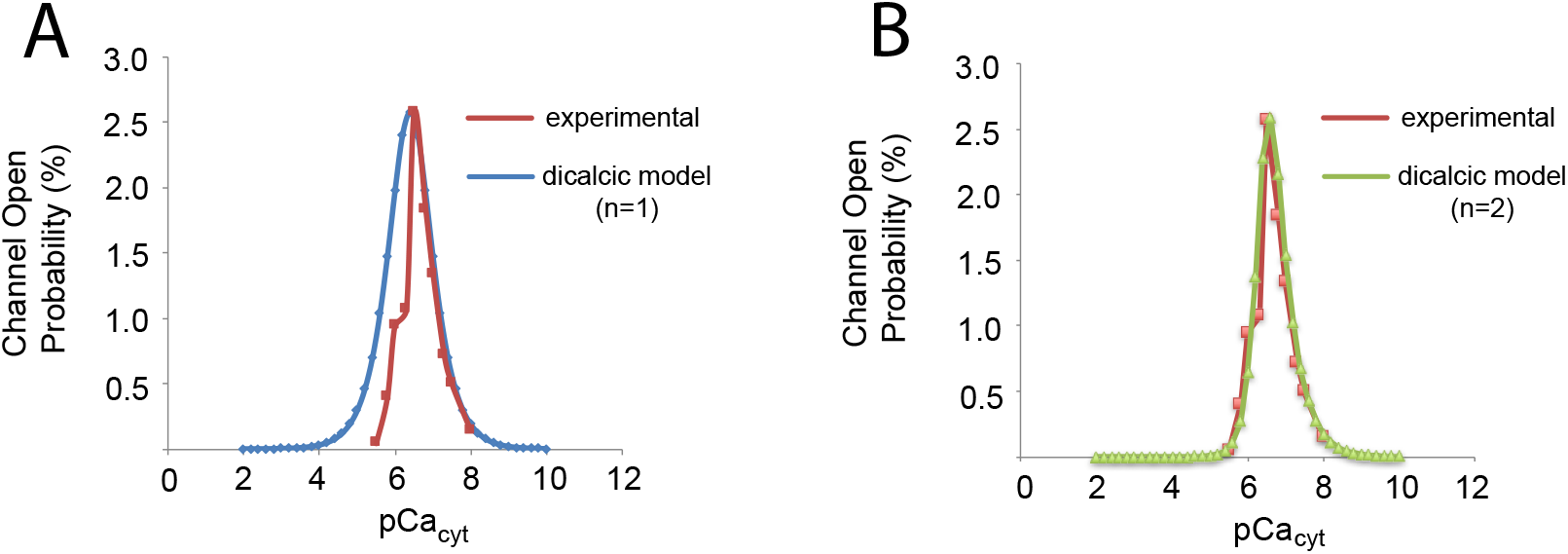
Channel Open Probability of IP_3_R in the presence of 2 *μ*M IP_3_ as a function of pCa_cyt_. The redrawn experimental data by Kaftan et al. [73] are outlined in red. Panel A shows the fit of the dicalcic model with a inhibiting cooperativity of 1. The discrepancy between model and experiment is clearly seen for high cytosolic Ca concentrations (low pCa_cyt_). In panel B the inhibiting cooperativity is 2, which results in a much better overall fit. The fits to all four experimental data sets by Kaftan et al. are presented in the Supporting Information S3 Text.

A possible explanation of the more abrupt IP_3_R inhibition at high Ca_cyt_ concentrations may be due to the binding of further Ca ions to the channel which then results in a conformational change inhibiting IP_3_R.

Fig 16 indicates the implementation of the dicalcic model into the overall cell model of calcium homeostasis. The parameters *k*_40_ and *k*_41_ play the respective roles of 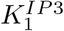 and 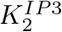; *k*_42_ and *k*_43_ represent 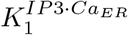 and 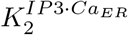, respectively. Parameters *k*_38_ and *k*_39_ are, respectively, the channel turnover number (*k*_2_ in Fig 14) and the *K*_*M*_ defined by Eq 25.

**Fig 16.**
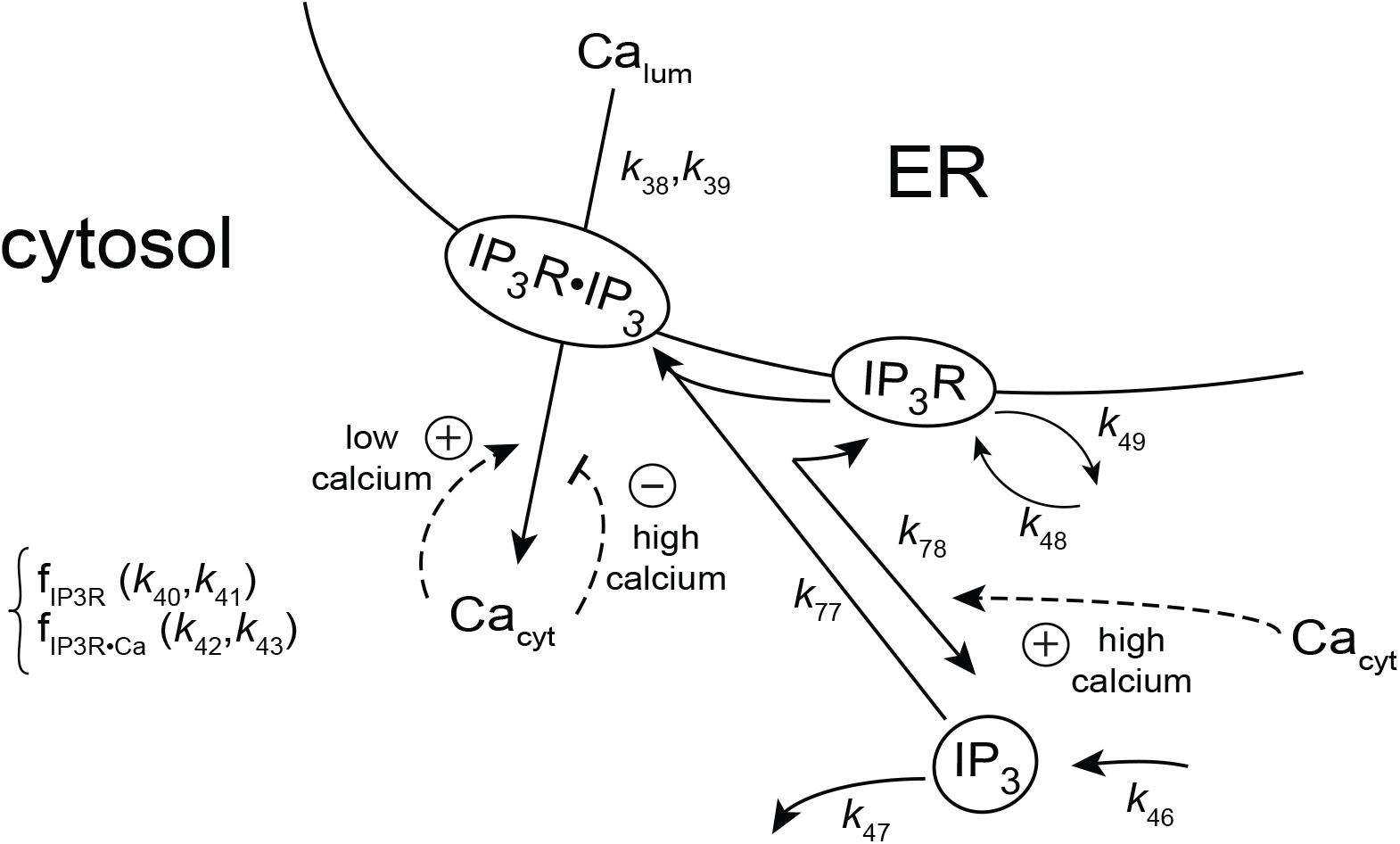
Incorporation of the dicalcic IP_3_R•IP_3_ channel activity (Fig 14) into the overall model. For a description of rate constant assignments, see text.

### Effect of cytosolic calcium on the dissociation of IP_3_R•IP_3_

We also investigated the possible outcomes when cytosolic calcium has an influence on the dissociation of IP_3_R•IP_3_, as outlined in Fig 16. We have modelled three scenarios how cytosolic calcium could influence the stability of IP_3_R•IP_3_:

i. considering a first-order kinetic influence of 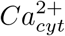 on the IP_3_R•IP_3_ dissociation by using the term *k*_78_·(*IP*_3_*R*•*IP*_3_)·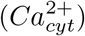
ii. considering the term 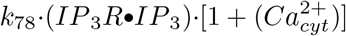, when only 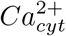 levels ≫ 1 *μ*M have a significant influence on IP_3_R•IP_3_ dissociation, or finally
iii. when 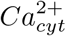 levels have no influence on IP_3_R•IP_3_ dissociation, i.e. using the dissociation term *k*_78_·(*IP*_3_*R*•*IP*_3_).

In order to test the three above conditions on the IP_3_R•IP_3_ transport rate *v*_*IP*3*R*_ we used a calcium inflow term *j*_1_=*k*_1_(*Ca*_*ext*_) into the cytosol which changed between 10 *μ*M/s and 4×10^4^ *μ*M/s. This caused a cytosolic calcium increase from 8.1×10^−4^ *μ*M to 4.8 *μ*M for all the three cases (i)-(iii) (Fig 17A). Interestingly, when 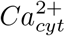 influences IP_3_R•IP_3_ dissociation by first-order kinetics (case (i)) the bell-shaped IP_3_R•IP_3_ transport rate disappears with high *v*_*IP*3*R*_ values at low *j*_1_ inflows (Fig 17B). Since experiments clearly show low IP_3_R•IP_3_ transport rates at low 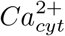 concentrations we conclude that an influence of low 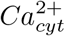 concentrations on IP_3_R•IP_3_ dissociation can be excluded. The results for case (ii) (influence of high 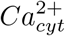 on IP_3_R•IP_3_ dissociation) and case (iii) (no influence of 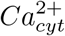 on IP_3_R•IP_3_ dissociation) are qualitatively similar (Fig 17B), i.e., an influence of high 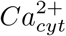 on IP_3_R•IP_3_ dissociation cannot be excluded.

**Fig 17.**
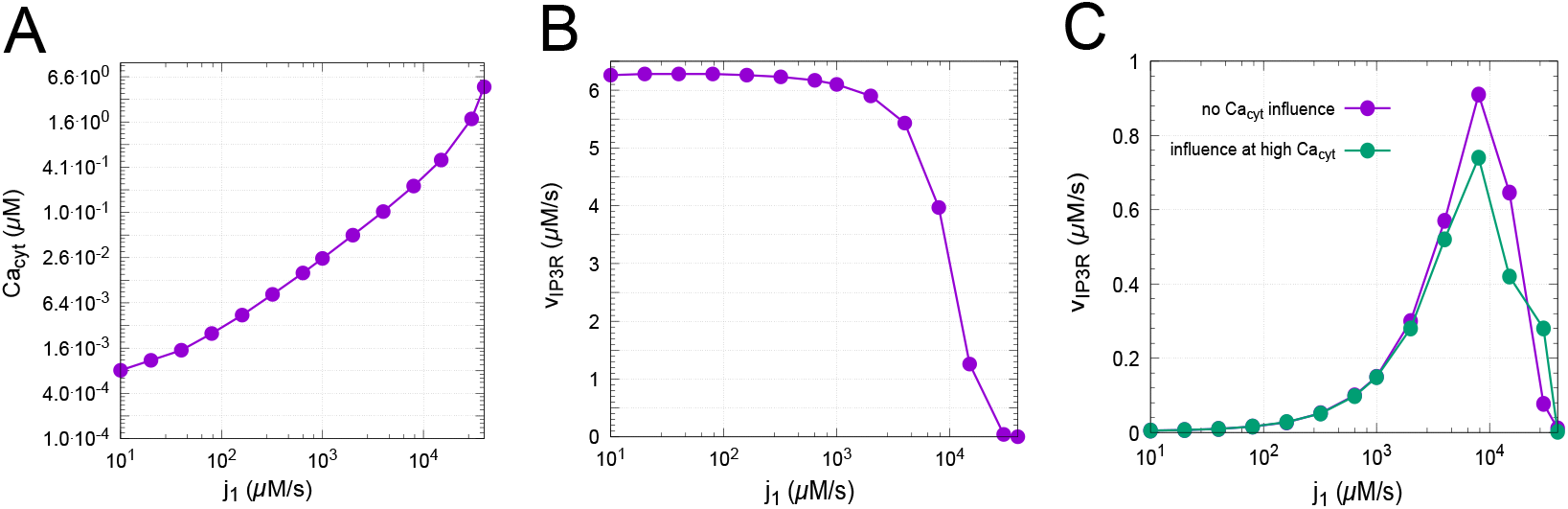
Influences of cytosolic calcium and cytosolic calcium inflow rates j_1_ on IP_3_R•IP_3_ dissociation and activity. Panel A: increase of cytosolic calcium concentration 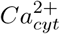 as a function of calcium inflow rate *j*_1_ into the cytosol (independent how 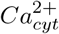 influences IP_3_R•IP_3_ dissociation). Panel B: IP_3_R•IP_3_ transport rate *v*_*IP*3*R*_ as a function of *j*_1_ when IP_3_R•IP_3_ dissociation is first-order with respect to 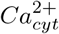; see (i) in text. Panel C: IP_3_R•IP_3_ transport rate *v*_*IP*3*R*_ when IP_3_R•IP_3_ dissociation is not dependent on 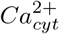 (purple curve, condition (iii) in text) or when high 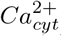 concentrations only influence IP_3_R•IP_3_ dissociation (green curve, condition (ii) in text). Details of these calculations are given in Supporting Information ‘S8 Program’.

### Oscillations

Although calcium oscillations [108–114] were not our primary focus in this research, we observed that the interaction between cytosolic calcium, PMCA/NCX, capacitative calcium entry and calcium-induced calcium release can lead to sustained oscillations. Remarkably, the period of these oscillations can vary from 2 seconds up to 30 hours! We found that the transport turnover numbers of SERCA (*k*_23_), IP_3_R (*k*_38_), the Ca set-point in the ER (defined by *k*_20_, *k*_21_), and the inflow rate of calcium into the cytosol (*k*_1_) are the parameters which affect the period most. For larger period lengths the oscillations are of the relaxation-type as shown in the build-up of calcium in the ER and its rapid release into the cytosol (Fig 18). The oscillations become high frequent with smaller amplitudes when *k*_23_ is increased (see ‘S9 Program’ for more details). In general, we found that an increased Ca inflow into the cytosol (up to a certain point) promotes (shorther period) oscillations, for example by the leak term *j*_1_=*k*_1_·*Ca*_*ext*_, or by a capacitative Ca entry at low Ca_ER_ levels. On the other hand, locking calcium concentrations in the ER to high values (by changing *k*_20_ and/or *k*_21_) leads to increased period lengths and eventually to the loss of oscillation. Concerning the IP_3_ channel oscillations, *j*_*IP*3*R*_, we sometimes observe a split peak as in Fig 18D. This occurs because of the bell-shaped behavior of the IP_3_R•IP_3_ Channel Open Probability (Fig 15). When the cytosolic calcium concentration increases above the pCa_cyt_ peak value (see Fig 15) and then decreases and passes the pCa_cyt_ peak again a double peak in *j*_*IP*3*R*_ is observed.

**Fig 18.**
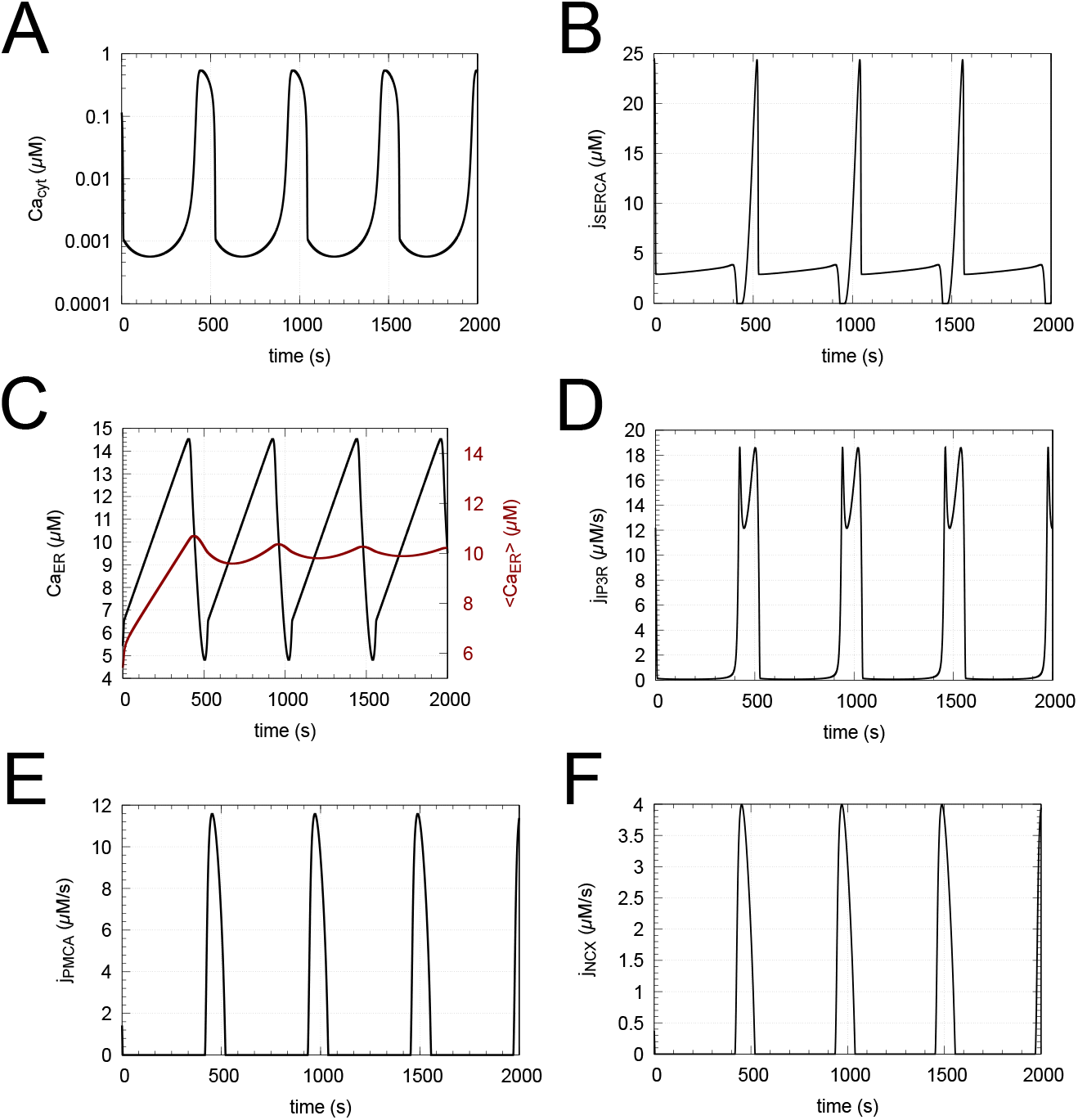
Example of sustained oscillations. The period in this case is 8.6 minutes. The red curve in panel C shows the average Ca_ER_ concentration which oscillates around its set-point (Eq 17), here set to 10 *μ*M. Also note the split peak of *j*_*IP*3*R*_ in panel D. Rate parameters, initial concentrations as well as binaries and the source file are given in ‘S9 Program’ together with additional calculations showing how parameter values can affect the oscillations with period lengths between a few seconds and up to 30 hours.

It should be noted that the oscillations shown in Fig 18, as well as the above referred 2s and 30h oscillations in ‘S9 Program’ are *independent* of IP_3_. In these calculations the levels of IP_3_R•IP_3_ and IP_3_ were kept constant by setting the rate constants *k*_77_ and *k*_78_ to zero. On the other hand, when the dissociation of IP_3_R•IP_3_ and its formation by IP_3_R and IP_3_ are taken into account, then different IP_3_ levels can increase or decrease period lengths by regulating the amount of IP_3_R•IP_3_ and thereby regulating the associated flux *j*_*IP*3*R*_ across the ER membrane (see *Influence of IP*_*3*_ in ‘S9 Program’).

Although the oscillations observed in our model could be classified as IP_3_ independent [111, 114], the kinetics and cooperativities of PMCA, NXC, leakage terms into the cytosol, and SOCC/STIM are additional factors which are expected to influence and contribute to the oscillations, besides IP_3_R•IP_3_ and IP_3_. It appears therefore interesting to investigate in further studies how the integration and their kinetics of the various inflows and outflows of cytosolic Ca^2+^ take part in the generation of these oscillations.

## Concluding remarks and future aspects

This work aimed at the development of a basic cellular model of Ca^2+^ homeostasis. A model like this poses the problem that certain cellular compounds/parts and their functions will certainly be neglected. In our case the role of mitochondria as an additional Ca store and regulator was not included, together with transporters such as the RyR’s, ARCC’s, VOCC’s along with sodium and potassium channels. Such neglects certainly limits the applicability of the model. On the other hand, we feel that through the model we have pointed at important regulatory features, such as the hysteretic behaviors of the Ca pumps and the enhanced cooperativity when the IP_3_R•IP_3_ channel closes at high Ca_cyt_ levels. The model’s astonishing oscillatory capability, which integrates Ca regulatory aspects of the model and is largely unexplored, appears to be an interesting aspect of further research with respect to both ultradian/high frequency [108, 110, 112] and possibly circadian [113] Ca oscillations. We also hope that the source files of the accompanying programs and their compiled binaries may be of interest as starting points for generating more improved models.

## Supporting information

**S1 Text Model details**. List of rate equations, abbreviations, and LSODE Y(i) assignments for the model Fig 3.

**S2 Text PMCA kinetic parameters for S2 Program and S3 Program**. Description of PMCA rate equations, cytosolic Ca^2+^ set point, and the estimation of PMCA V_max_, K_*M*_ and turnover number.

**S3 Text Dicalcic model**. Derivation of Eq 22 with n=1 and n=2, and fits to the experimental data by Kaftan et al. (Fig 2 in Ref. [73]).

**S1 Program Cytosolic calcium homeostasis in a minimal model**. Fortran source file and executables showing results of Fig 2.

**S2 Program Modeling the application of ionophore A23187 on erythrocytes**. Rate equations, parameters and calculations for Fig 5B.

**S3 Program Modeling cytosolic calcium regulation by PMCA in bovine endothelial cells**. Rate parameters and calculations for Figs 6B and 6C.

**S4 Program Modeling cytosolic calcium regulation by PMCA and NCX in bovine endothelial cells**. Rate equations, parameters, initial concentrations, and calculations for Figs 7C and 7D.

**S5 Program Modeling cytosolic calcium regulation by PMCA and NCX in bovine endothelial cells**. Rate equations, parameters, initial concentrations, and calculations for Fig 8B.

**S6 Program Analysis of Ca**^**2+**^**-leakage rates from the ER**. Determination of first-order rate constant and Michaelis-Menten as well as hysteretic parameters from the experimental results by Luik et al. [74] and Camello et al. [10], respectively.

**S7 Program Calculations leading to Figs 10-12**. Calculated capacitative Ca^2+^ inflow rates with concentration dependent cooperativity.

**S8 Program Influences of cytosolic calcium and cytosolic inflow rates** *j*_1_ **on IP**_**3**_**R**•**IP**_**3**_ **dissociation and activity**. The supporting information shows the details leading to Fig 17.

**S9 Program Oscillatory responses of the model**. Calculations showing the results of Fig 18 and parameter influences on the period.

## Acknowledgments

We thank Tormod Drengstig and Kristian Thorsen for comments during the early stages of this project.

## Notes

### Competing Interest Statement

The authors have declared no competing interest.

